# Systematic classification of phage receptor-binding proteins predicts surface glycopolymer structure in *Staphylococcus* pathogens

**DOI:** 10.1101/2024.03.04.583386

**Authors:** Janes Krusche, Christian Beck, Esther Lehmann, David Gerlach, Christiane Wolz, Andreas Peschel

## Abstract

Wall teichoic acids (WTAs) are major surface polymers of staphylococcal pathogens and commensals, whose variable structure governs interaction with host receptors, immunoglobulins, and bacteriophages. The ribitol phosphate (RboP) WTA type contributes to virulence, for instance in *Staphylococcus aureus*, but we lack comprehensive knowledge of WTA types and cognate phages.

We developed a computational pipeline to identify the receptor-binding proteins (RBPs) in 335 Staphylococcus phage genomes, yielding multiple distinct RBP clusters. Notably, many phages had two separate RBPs with in part different WTA preferences. RBP representatives differed in specificity for RboP WTA glycosylation types, recapitulating the specificity of the corresponding phage. Based on these results, we created a publicly available bioinformatic tool to predict phage host specificity based on RBP similarity.

The RboP WTA specific Φ13-RBP also revealed that the presence of RboP WTA on non-aureus staphylococci is more common than previously thought. Our approach facilitates the characterization of opportunistic Staphylococcus pathogens according to WTA types, which has major implications for phage-mediated interspecies horizontal gene transfer and future phage therapies.

## Introduction

Infections with antibiotic resistant bacterial pathogens, including *Staphylococcus aureus,* threaten human health worldwide. In 2019, *S. aureus* infections alone led to more than 1 million deaths (Antimicrobial Resistance Collaborators, 2022b), more than 700,000 of which were associated with antimicrobial resistance (Antimicrobial Resistance Collaborators, 2022a). The urgent need for treatment strategies alternative to antibiotics has revived the interest into bacteriophages, viruses that infect and kill bacteria. In addition to the therapeutic potential of phages against infections with antibiotic resistant bacteria, the narrow host range of phages decreases unwanted effects on other, potentially beneficial bacteria during treatment (Mu et al., 2021).

All currently known *S. aureus* phages belong to the order caudovirales, consisting of three morphologically separate groups (Xia & Wolz, 2014). Podoviruses are small phages with a very short, non-contractile tail and a limited host range. Siphoviruses possess long, non-contractile tails and they can alternate between the lysogenic and lytic lifecycle. Accordingly, siphoviruses can be found in the form of prophages in many *S. aureus* genomes (Ingmer et al., 2019; Xia & Wolz, 2014). Myoviruses have the largest genome of the three groups, carrying genes for a contractile tail as well as many accessory genes such as tRNAs and nucleases (O’Flaherty et al., 2004). *S. aureus* strains encode only a limited number of phage defense systems, the most prevalent being restriction-modification and abortive infection systems (Jurado et al., 2022), whereas Clustered Regularly Interspaced Short Palindromic Repeats (CRISPR)-Cas systems can only be found in 2.9% of *S. aureus* genomes (Mikkelsen et al., 2023). Furthermore, alteration of wall teichoic acid (WTA) polymers, can protect from phage infection. While phages of other bacterial groups often use different receptors on host cells such as lipopolysaccharides or membrane proteins (Leprince & Mahillon, 2023), phages of staphylococci appear to use only one receptor, the peptidoglycan-linked WTA polymer consisting of ribitol-phosphate (RboP) repeats (Weidenmaier & Peschel, 2008; Xia et al., 2011).

The RboP WTA backbone of *S. aureus* is decorated with N-acetylglucosamine (GlcNAc) in various conformations depending on the presence or absence of specific genome-encoded glycosyltransferases (van Dalen et al., 2020). There are currently three RboP WTA glycosyltransferases known in *S. aureus*. The main glycosyltransferase, TarS, mediates attachment of β-1,4-GlcNAc. It is present in the majority of *S. aureus* strains, while the accessory glycosyltransferases TarM (responsible for WTA α-1,4-GlcNAc modification) and TarP (WTA β-1,3-GlcNAc modification) can only be found in 37% and 7% of *S. aureus* strains, respectively (Brown et al., 2012; Gerlach et al., 2018; Gerlach et al., 2022; Tamminga et al., 2022; Xia et al., 2010). Changes in WTA glycosylation have been shown to impact phage binding and resistance to some phages (Li et al., 2015; Winstel et al., 2014; Yang et al., 2023).

The relationship of *S. aureus* phages and phages of non-*aureus* staphylococci (NAS) such as *Staphylococcus epidermidis* has remained unclear. While the cell wall of *S. aureus* is decorated with RboP WTA, most NAS carry WTA consisting of glycerol-phosphate repeats (GroP) with various sugar side chains such as glucose or GlcNAc (Beck et al., 2024; Endl et al., 1983). Due to the fundamental difference in WTA backbone composition between *S. aureus* and NAS, most phages of *S. aureus* are not able to bind or infect NAS such as *S. epidermidis*, and vice versa. This division limits horizontal gene transfer by phages (Winstel et al., 2013), and it is unclear how resistance genes such as the methicillin resistance gene *mecA,* encoded on the staphylococcal chromosomal cassette *mec* (*SCCmec*), have been transferred from NAS to *S. aureus* in the past (Rolo et al., 2017). Nevertheless, phage transduction is believed to be the major mechanism of horizontal gene transfer in the genus *Staphylococcus*.

Phage adsorption, the initial step of phage infection and transduction, depends on successful attachment of the phage to its host and thus on the specificity of the phage receptor-binding proteins (RBPs). Adsorption is often established in a three-step process (Bertozzi Silva et al., 2016). First, contact between the phage and its host happens via random diffusion, leading to reversible binding of one RBP to a receptor on the host surface. This initial step is followed by binding of a second RBP that irreversibly binds to a second receptor (Garen & Puck, 1951), facilitating a conformational change, which ultimately results in injection of viral DNA into the host. While this infection process has been shown for the *E. coli* phage T1 (Bertozzi Silva et al., 2016), only little is known about the adsorption mechanism to *S. aureus.* Furthermore, the RBPs of *S. aureus* phages have not been identified except for a few model phages. The RBP structures of the siphovirus Φ11 and the closely related Φ80α have been determined by crystallization (Kizziah et al., 2020; Koc et al., 2016), as well as the full baseplates with integrated RBPs of podovirus ΦP68 (Hrebik et al., 2019). For myoviruses, the binding behaviors of the two RBPs of ΦSA012 have been found to be glycosylation-dependent, although a clear binding pattern remains to be elucidated (Takeuchi et al., 2016).

To prevent and treat multi-resistant staphylococcal infections by future phage therapies, it is vital to understand the transduction-based dissemination of virulence and resistance genes, as well as the mechanisms by which phages infect and kill *S. aureus*. Here we combined computational and experimental approaches to identify several novel RBPs in most of the known *S. aureus* phages. Many of these phages appear to encode two different RBPs, some of which were capable of reversible binding, indicating that some *S. aureus* phages follow the classical three-step infection pattern of contact by diffusion, then reversible, followed by irreversible attachment. Most RBPs of *S. aureus* phages can be grouped in several distinct clusters with high overall similarity. We show that RBP binding is WTA dependent, but that the glycosylation-dependent binding specificity varies from one RBP cluster to another. The binding of recombinant RBPs matches the behavior of the corresponding phages, substantiating the notion that the identified RBPs govern phage host range. By combining phylogenetic and binding specificity analysis of RBPs, we can predict the binding pattern of nearly all known *S. aureus* phages. To this end, we developed an easy-to use tool called **Ph**age **A**ureus **R**BP **I**dentification **S**ystem (PhARIS), enabling the user to predict RBPs from phage genomes and thus assess binding specificity of any *S. aureus* phage based on its genome sequence.

## Results

### RBPs of *S. aureus* phages can be grouped into separate clusters according to amino acid sequence similarity

Phage infection requires adsorption to the host cell but the phage determinants governing host-specific binding have remained only superficially understood. To elucidate the receptor specificities of *S. aureus* phages, all published information about RBP loci of the *S. aureus* siphoviruses Φ80α and Φ11, myovirus ΦSA012, and podoviruses ΦP68 and ΦS24-1, which encode partially characterized RBPs (Hrebik et al., 2019; Kizziah et al., 2020; Koc et al., 2016; Takeuchi et al., 2016; Uchiyama et al., 2017) was collected. Additionally, new putative RBPs of siphoviruses Φ12 and Φ13 and of multiple *S. aureus*-infecting myoviruses were discovered via HHpred, BLAST, AlphaFold2 and by comparison of gene locations, based on the fact that structural genes including RBP genes often show synteny in phage genomes (Jumper et al., 2021; Soding et al., 2005). The resulting curated list of RBPs was used for searching further RBP homologs in a database of 479 available *Staphylococcus* phage genomes, which was created by extracting all *Staphylococcus* phage genomes from the INPHARED phage database (April 1^st^, 2023) (Cook et al., 2021). 335 of 479 phage genomes (69.94%) encoded proteins that matched at least one of the RBPs with more than 65% identity and 65% overlap. RBP genes of 144 phages could not be assigned, most of which were phages of NAS, indicating strong differences between phage RBPs of *S. aureus* and NAS. According to the NCBI database, only 10 of the remaining 144 phages have *S. aureus* as their host, 7 of which are giant viruses whose RBPs appear to be unrelated to those of other phage groups.

Following the identification of RBP genes, multiple alignment of protein sequences was performed with Clustal Omega (Sievers et al., 2011). The resulting phylogenetic tree (Figure 1b) revealed the existence of several distinct clusters of potential *S. aureus-*specific RBPs with very high intra-cluster similarity, and very low similarity to the next-closest cluster. The cluster of RBPs from Triaviruses was found to have the highest intrinsic relatedness, with a 97.8% identity of the query (Φ12-RBP) to the most distant match of the cluster. The next closest match outside of this cluster was found in the Azeredovirinae RBP cluster, with an identity to the Φ12-RBP of 50.9%.

**Figure 1:**
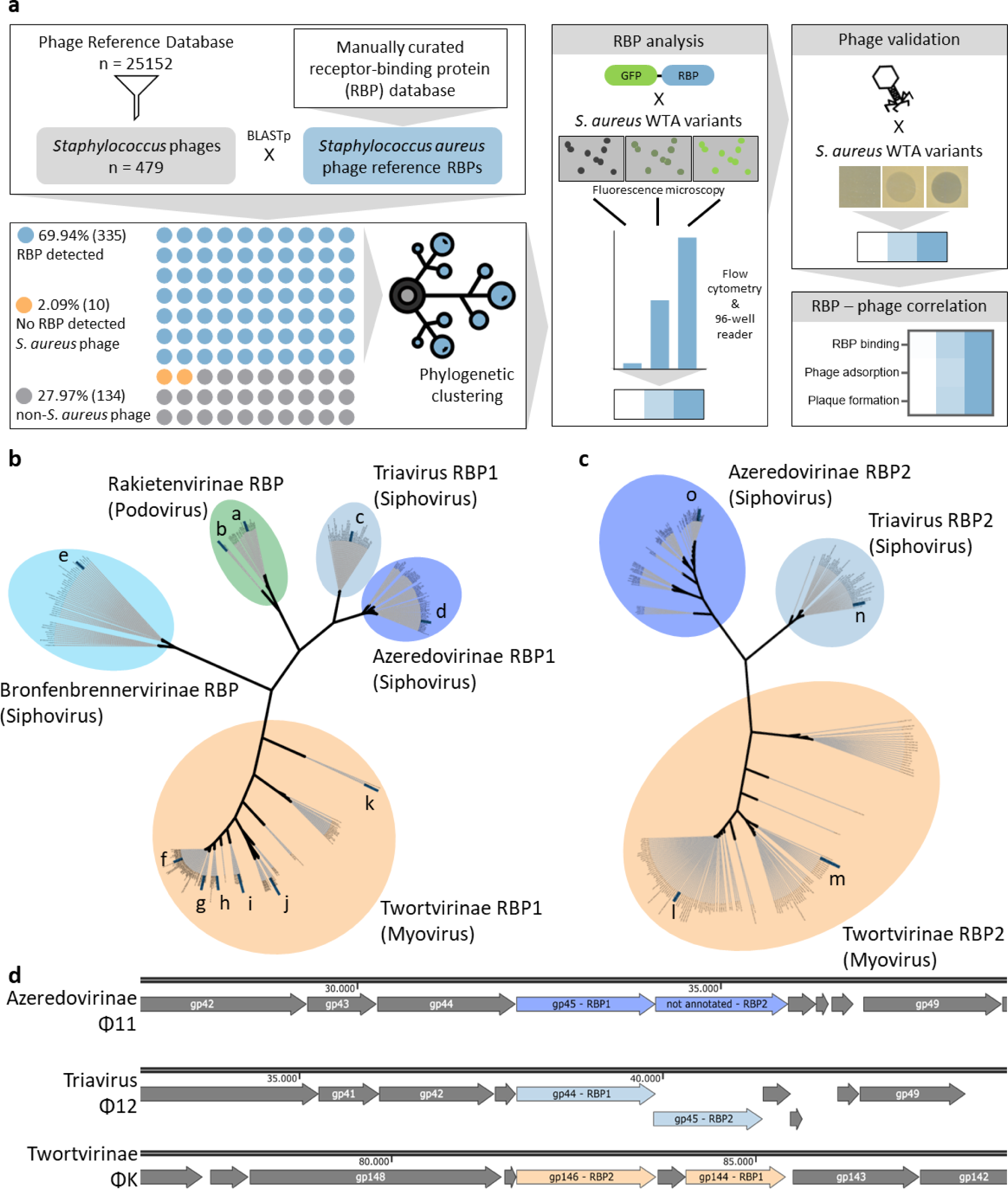
*S. aureus* phages encode one or two RBPs that show phylogenetic clustering with high intra-cluster homology. **a,** Schematic representation of the bioinformatic (left) and wet-lab (right) workflow that was used to classify and characterize *S. aureus* RBPs. A reference RBP panel, consisting of multiple typical representative RBPs, was aligned with a database of 479 *Staphylococcus* phage genomes from the INPHARED database, resulting in 335 phages with at least one RBP match (69.94%). In 10 *S. aureus* phage genomes (seven of those were giant viruses), no RBP could be detected (2.09%). The remaining 134 genomes belonged to non-*S. aureus* phages (27.97%) where no RBP could be detected. The resulting collection of *S. aureus* RBPs was clustered according to amino acid sequence homology. Per cluster, one or more green fluorescent protein (GFP)-RBP fusion-proteins were designed, overexpressed, and analyzed for binding specificity. RBP binding results were compared to experiments of phage infection and adsorption. **b,** *S. aureus* phage RBPs (RBP1) cluster according to their amino acid sequence. Podovirus RBPs could be separated into two subclusters (a: ΦP68-like and b: ΦCSA13-like), while siphovirus RBPs could be clustered into three different subclusters (c: Φ12-like (Triavirus), d: Φ11-like (Azeredovirinae) and e: Φ13-like (Bronfenbrennervirinae)). Myovirus RBPs could be separated into six different subclusters (f: ΦK-like, g: ΦStab20-like, h: ΦSA012-like, i: ΦPG2021-10-like, j: ΦPG2021-17-like, k: ΦBS1-like). **c,** A second RBP (RBP2) was only detected in myoviruses (l: ΦK-like, m: ΦPG2021-10-like) and two clusters of siphoviruses (n: Φ12-like (Triavirus), o: Φ11-like (Azeredovirinae)). A second RBP was not found in Rakietenvirinae (podoviruses) and Bronfenbrennervirinae. **d,** Genomic location of RBP1 and RBP2 in one representative of each double-RBP containing group (Azeredovirinae, Triaviruses and Twortvirinae).

Based on the similarity of the RBPs, the *S. aureus* phage RBPs could be assigned to five main clusters with several subclusters. The podovirus RBPs clustered into two distinct subclusters represented by the RBPs of ΦP68 (Figure 1b, a) and ΦCSA13 (Figure 1b, b), as described previously (Uchiyama et al., 2017). The siphovirus RBPs clustered into three clusters: those of Azeredovirinae (represented by Φ11 (Figure 1b, d)), of Triaviruses (represented by Φ12 (Figure 1b, c)), and of the more distant Bronfenbrennervirinae (represented by Φ13 (Figure 1b, e)). The vast majority of currently known *S. aureus* podo- and siphoviruses (99%) could be assigned to one of these four clusters. Myovirus RBPs displayed higher heterogeneity compared to podo- and siphovirus RBPs. For this reason, a 65%-identity cutoff was used to filter out RBPs of NAS-infecting myoviruses. The remaining *S. aureus*-specific myoviruses could be clustered into six subclusters.

In the azeredovirus Φ80α, a close relative of Φ11, the existence of a second RBP (RBP2) on its baseplate has previously been hypothesized but a functional characterization of this hypothetical second RBP has yet to be performed (Kizziah et al., 2020). In the myovirus ΦSA012, functional host adsorption has been demonstrated for two RBPs, RBP1 and RBP2. While these two RBPs were found to have different binding properties during phage infection, the exact role of each RBP has remained unclear (Takeuchi et al., 2016). As the presence of two distinct RBPs may be a more general feature of *Staphylococcus*-specific phages, we set out to identify potential secondary RBPs in the entire *Staphylococcus* phage genome database. The RBP2 sequences were analyzed in a similar way as those of the first RBPs (Figure 1c). In the majority of podoviridae as well as the Bronfenbrennervirinae cluster of siphoviruses, no RBP2 could be detected. In contrast, the siphovirus type Triavirus carries a distinct, putative RBP2, which is encoded directly downstream of RBP1 (Figure 1d). Due to the organization of phage genomes into functional modules, genes encoding similar functions are often found in close proximity. In fact, we found the putative RBP2 genes of Azeredovirinae, Triaviruses, and Twortvirinae (myoviruses) to be located directly next to those of RBP1 (Figure 1d). Based on this observation, we could predict a putative RBP2 in Φ11, Φ12, and ΦK (Figure 1c). Myoviruses other than ΦK likely also carry RBP2 (Figure 1c, l & m).

Overall, these results show a high intra-cluster conservation across *S. aureus* phage RBPs, indicating that the evolutionary pressure to change RBP architecture is low. The conserved nature of RBPs one or two could be due to the highly conserved structure of *S. aureus* WTA, prompting us to investigate the binding behavior of these proteins to the WTA of the host.

### RBP1 of ΦK shows irreversible binding to WTA while RBP2 binds reversibly to WTA, indicating the classical three-step binding pattern for *S. aureus* phages

To analyze the binding capability of the *in-silico* detected putative RBPs, interaction of recombinant RBP proteins with different WTA variants, the presumed RBP ligands, was investigated. To this end, a selection of RBPs fused to green fluorescent protein (GFP) (See Table S3) were produced in *Escherichia coli*, purified, and tested for binding to various WTA mutants of *S. aureus* USA300 JE2, a methicillin-resistant strain that carries both TarM and TarS (Diep et al., 2006), as well as to *S. epidermidis* 1457 expressing GroP WTA. The fluorescence of the bacteria-adsorbed proteins was quantified via fluorescence microscopy and flow cytometry. Here, we focused on the myovirus ΦK due to its great potential as an agent in phage therapy (Atshan et al., 2023; Lehman et al., 2019). RBP1 of ΦK was found to bind to *S. aureus,* irrespective of the glycosylation state of RboP WTA, and to *S. epidermidis*, but it was not able to bind the WTA-deficient *S. aureus* Δ*tagO* mutant (Figure 2a & 2b) (D’Elia et al., 2009). Specifically, RBP1 of ΦK showed high affinity to *S. aureus* WT, Δ*tarM,* and Δ*tarS,* all carrying glycosylated RboP WTA, as well as to the GroP WTA expressing *S. epidermidis*, and intermediary affinity to glycosylation-deficient *S. aureus* Δ*tarM* Δ*tarS.* Interestingly, the native phage K showed strong binding affinity to, and plaque formation on, *S. aureus*, independent of the glycosylation state of the RboP WTA. Weaker affinity was observed for S. *epidermidis* 1457 (Figure S3). To elucidate this discrepancy, the influence of the putative second RBP on the host affinity of ΦK was assessed.

**Figure 2:**
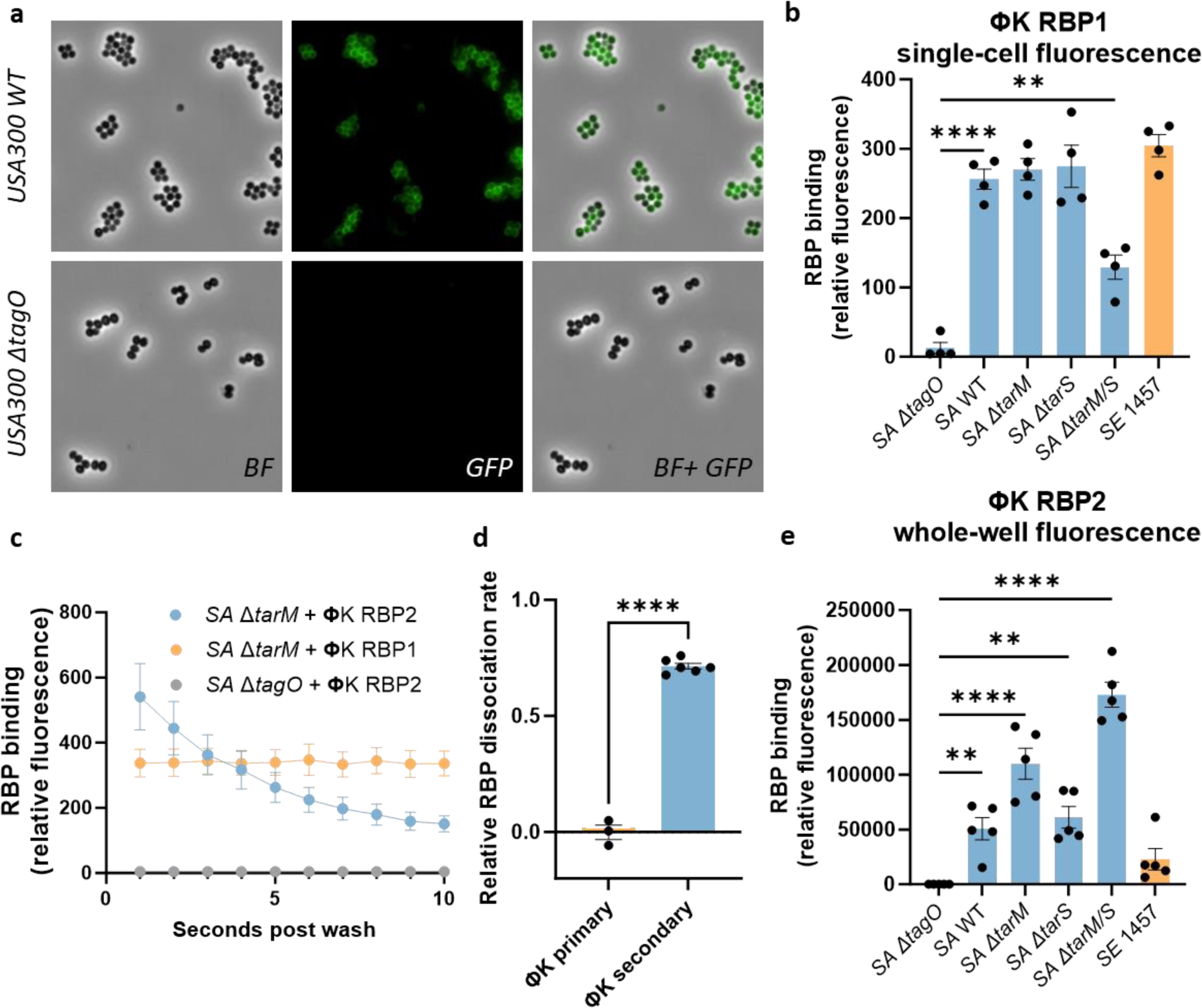
ΦK carries two RBPs for reversible and irreversible binding, both shaping the infection behavior of the phage. Fluorescent GFP-RBP constructs were incubated with various *S. aureus* USA300 JE2 WTA glycosyltransferase mutants, as well as a WTA-deficient control mutant (*S. aureus* USA300 JE2 Δ*tagO*) and with GroP WTA containing *S. epidermidis* 1457. Fluorescence of the ΦK RBP1-bound bacteria was assessed via **a** microscopy and **b** flow cytometry. Quantification was achieved by measuring the median fluorescence intensity. RBP2 binding of ΦK was measured over time via **c** flow cytometry and in a **e** fluorescence reader in 96-well format. **d,** The dissociation rate of the RBPs was calculated as relative decrease in fluorescence between 1 and 10 seconds after washing. Data for all tested RBPs can be found in Figure S1 & 2. Statistical analysis was done via **b, e** ordinary one-way ANOVA or **d** t-test, and multiple comparisons were performed between the WTA-negative *S. aureus* Δ*tagO* strain and the differently glycosylated strains, **P < 0.01, ****P < 0.0001.

RBP2 of ΦK had a pronounced capacity to bind to all tested *S. aureus* strains except for the WTA-deficient Δ*tagO* mutant, which confirms its role as a WTA-binding RBP. The affinity of ΦK RBP2 to the *S. aureus* Δ*tarM* and Δ*tarM* Δ*tarS* mutants was two- to three-fold higher compared to the parental USA300 JE2 WT strain and the Δ*tarS* mutant (Figure S2). Interestingly, RBP2 of ΦK bound to *S. aureus* but dissociated quickly after washing of the RBP-bound bacterial cells, which was visible as a time-dependent decrease of fluorescence in the flow cytometer (Figure 2c & 2d). This behavior was clearly not observed for RBP1 of ΦK. This finding indicates that while RBP1 binds irreversibly, RBP2 can bind to WTA only in a reversible manner.

As we suspected that reversibly bound RBPs could get lost during flow cytometry analysis after washing of the bacteria, leading to lower binding rates, we developed an additional assay based on fluorescence in microtiter plate wells to allow for quantification of RBP binding. In this microtiter plate assay, the bacteria were mixed with RBP2 and washed once, after which the fluorescence of the whole well was measured. Under these conditions, most fluorescent proteins initially associated with the bacteria should remain in the samples, irrespective of potential subsequent dissociation. In this assay, the RBP2 of ΦK was found to have similar affinities for the various *S. aureus* and *S. epidermidis* test strains compared to the flow cytometry assay. The RBP2 of ΦK adsorbed most effectively to USA300 Δ*tarM* Δ*tarS* with unglycosylated RboP WTA but was also able to bind every other *S. aureus* strain carrying RboP WTA (Figure 2e). The strong binding of ΦK to all *S. aureus* strains, including the Δ*tarM* Δ*tarS* deficient strain, can thus be explained by strong cooperative binding of both RBPs. While RBP1 preferentially binds to glycosylated WTA in a stable fashion, the reversibly binding RBP2 prefers unglycosylated RboP WTA. By contrast, binding of ΦK to GroP WTA-bearing *S. epidermidis* appears to be largely mediated by RBP1. We hypothesize that the weak contribution of RBP2 to *S. epidermidis* binding may be the reason for the comparatively low binding efficiency and thus reduced infection of *S. epidermidis* by ΦK (Figure S3).

We expanded the principles observed for the mechanisms of RBP receptor interactions to other double RBP carrying phages. Using computational and functional approaches, various novel RBPs of *S. aureus*-specific phages were found, including the second RBPs of myoviruses ΦSA012 and ΦStab20 as well as siphoviruses Φ11 and Φ12 (Figure 3).

**Figure 3:**
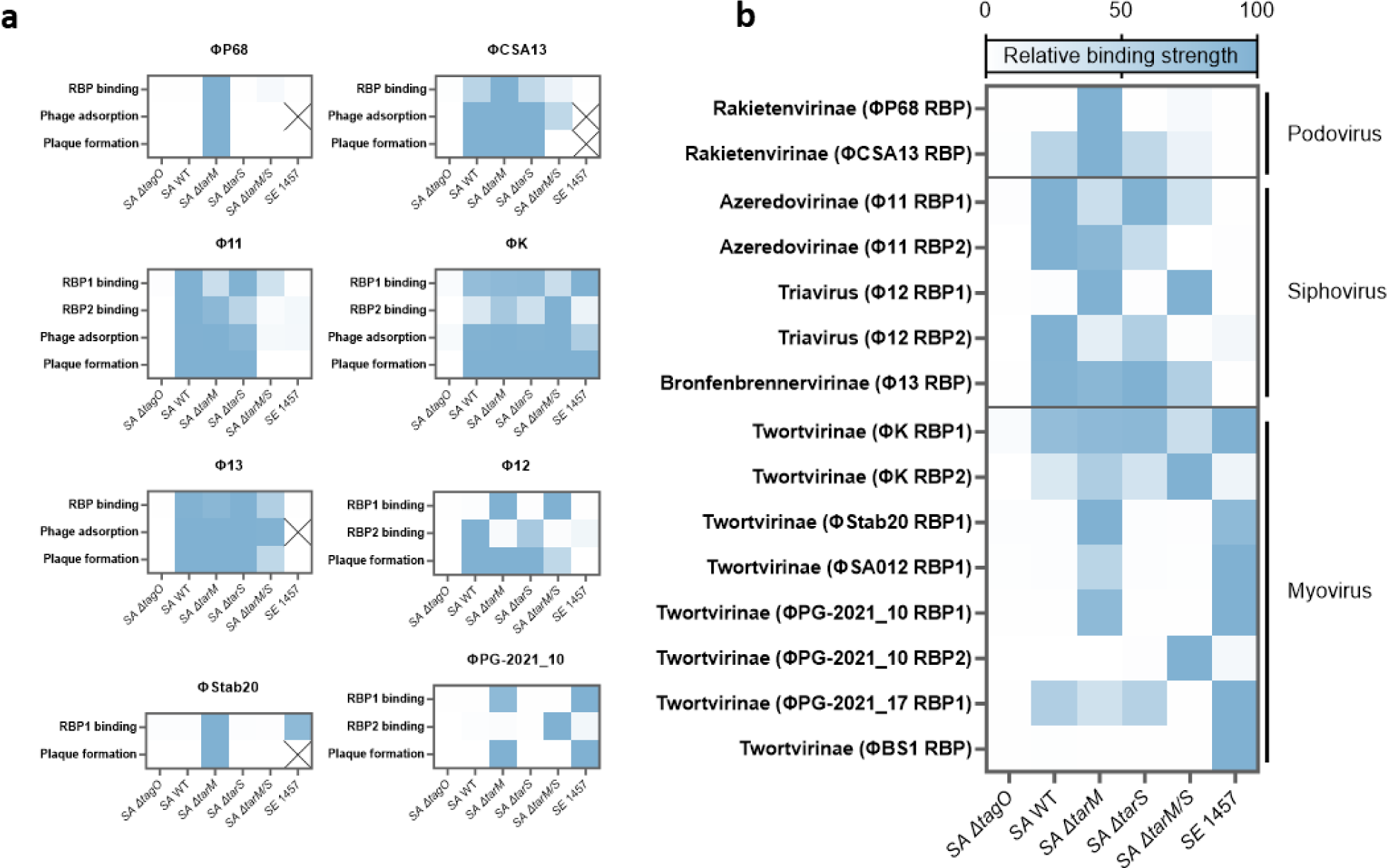
RBP binding matches with behavior of phages. **a,** Phage adsorption and plaque formation of various phages in comparison with RBP binding. The host range of podoviruses ΦP68 and ΦCSA13, as well as siphovirus Φ13 can be explained by one RBP alone. The siphoviruses Φ11 and Φ12, as well as the myoviruses ΦK, ΦStab20 (same subcluster as the RBP2 of ΦPG2021-10) and ΦPG2021-10 carry a second RBP (RBP2) that contributes to the phage host range. **b,** WTA-glycosylation dependent binding pattern of *S. aureus* phage RBPs. Distinct RBP subclusters show different WTA binding specificities. None of the RBPs could bind to the WTA-deficient Δ*tagO* and only myovirus RBPs could bind to both, the GroP WTA of *S. epidermidis* as well as the RboP WTA of *S. aureus*. For quantification of RBP adsorption, values were normalized to the highest fluorescence of each RBP. Plaque formation was classified into strong lysis (100%, blue), weak lysis (50%, light blue) and no lysis (0%, white). Numerical data can be found in Figure S5.

### RBP binding patterns match phage behavior and can predict WTA-dependent host range of characterized and uncharacterized phages

Phage-*S. aureus* interactions have previously been studied using only a small set of model phages. The receptor specificity of most *S. aureus* phages, with regard, for instance, to the specific WTA glycosylation pattern, has remained superficially understood (Gerlach et al., 2018; Li et al., 2015; Xia et al., 2010). We assessed one or more representative RBPs of each of the RBP clusters (Figure 1) for their WTA binding properties to elucidate the molecular basis of *S. aureus* phage recognition.

The cluster of podoviruses, which carry only one RBP, can be separated into two subclusters. ΦP68, a representative of the first subcluster, infects exclusively Δ*tarM* mutants with only β-1,4-GlcNAc glycosylated WTA, as shown previously (Li et al., 2015). Similarly, the RBP of ΦP68 (Figure 3a and b) only showed strong adsorption to the Δ*tarM* mutant of USA300 JE2 and minimal binding to the unglycosylated Δ*tarM* Δ*tarS* strain, although this low binding did not correspond to the adsorption or infection pattern of the native phage (Figure 3a).

As surrogate for the podovirus ΦCSA13, whose RBP was chosen as reference RBP for this second subcluster (Figure 1a, b), published data of the closely related phage ΦS24-1 was used for whole phage assays, since its RBP also clusters in the second subcluster (92.5% sequence identity to the RBP of ΦCSA13). ΦS24-1 has been unable to infect *S. aureus* RN4220 Δ*tarM* Δ*tarS* (lacking WTA glycosylation) and Δ*tagO* (lacking WTA), but strongly lysed the wild type RN4220 and glycosyltransferase mutants Δ*tarM* and Δ*tarS* (Uchiyama et al., 2017). In adsorption experiments, S24-1 did not bind to Δ*tagO* but was able to bind the Δ*tarM* Δ*tar*S mutant to some extent (Figure 3a). The RBP of ΦCSA13 showed the same binding pattern, with no binding to the Δ*tagO* mutant and some, albeit very low, binding to the Δ*tarM* Δ*tarS* mutant.

The binding pattern of the siphovirus Φ11 RBP1 (Figure 3a) differed in our experiment from previously described data (Li et al., 2016). In the present work, the background fluorescence was minimized by using GFP fusion proteins instead of biotin-labelled RBPs, which allowed more sensitive measurements, revealing a difference in binding between Δ*tagO* and Δ*tarM* Δ*tarS*. *S. aureus* Δ*tarM* Δ*tarS* was bound strongly by the Φ11 RBP1, while the WTA-deficient strain could not be bound (Figure 3a). This finding shows that RBP1 of Φ11 is able to bind to unglycosylated WTA. However, Φ11 was neither able to bind to nor infect the Δ*tarM* Δ*tarS* strains (Figure 3a, Figure S5). To analyze the discrepancy between binding capacity of the RBP and the full phage, the putative RBP2 of Φ11 was investigated. RBP2 of Φ11 did not show any binding in flow cytometry, which could be due to the short-lived binding, followed by quick dissociation. Because of this instability, the above-mentioned microtiter fluorescence reader assay was used to analyze binding of RBP2. Compared to the negative Δ*tagO* control, we detected strong binding of Φ11 RBP2 to USA300 WT, Δ*tarM* and Δ*tarS*, while we observed no binding to the *S. aureus* Δ*tarM* Δ*tarS* mutant or to *S. epidermidis* 1457 (Figure 3, Figure S3). This pattern indicates that Φ11-like siphoviruses probably follow the three-step binding sequence as described for many other phages. Φ11-like siphoviruses make initial contact with their host via random diffusion, followed by reversible binding of RBP2 to the WTA, whereafter RBP1 binds irreversibly. The inability of these phages to infect the Δ*tarM* Δ*tarS* mutant lies in the lack of affinity of RBP2 to unglycosylated WTA.

RBP1 of the Triaviruses Φ3A and Φ47 has 99.8% and 100% identity to RBP1 of Φ12. Interestingly, the binding of the Φ12 RBP1 did not match the behavior of those phages (Figure 3a). An RBP2 can be found in all Triaviruses and is able to account for the differences in binding by RBP1 and the full phage (Figure 3a). The phages of this cluster were found to infect *S. aureus* irrespective of glycosylation, although the plating efficiency was slightly reduced for the Δ*tarM* Δ*tarS* mutant compared to the wild type (Figure S3). While the RBP1 of Φ12 only bound to the Δ*tarM* and Δ*tarM* Δ*tarS* mutant, RBP2 strongly bound to the WT and Δ*tarS* mutant. This pattern could indicate that both RBPs of Φ12 contribute equally to the binding mode of the phage.

Siphovirus Φ13 was found to bind strongly to the *S. aureus* wild type as well as to its glycosyltransferase single mutants, and showed weaker, albeit still robust, binding to the double-glycosyltransferase deficient Δ*tarM* Δ*tarS*. As a typical representative of the Bronfenbrennervirinae, Φ13 encodes only one putative RBP. In adsorption experiments, Φ13 bound to *S. aureus* test strains in a similar way as its RBP, although differences between Δ*tarM* Δ*tarS* and the other strains were only weakly noticeable, which was probably due to the overall high adsorption capacity (Figure 3, Figure S5). In the spot assay, we observed lysis zones on *S. aureus* RN4220 WT, RN442 Δ*tarM*, and RN4220 Δ*tarS*, and a more opaque zone on RN4220 Δ*tarM* Δ*tarS*. This indicates that infection of Φ13 only requires binding of its single RBP, and that this RBP can bind to RboP WTA irrespective of its glycosylation pattern.

As described above, myovirus ΦK carries two RBPs, both of which are able to bind to all strains of *S. aureus* except the Δ*tagO* mutant. Interestingly, *S. epidermidis* 1457 was only bound by RBP1, but not by RBP2 of ΦK. This difference could explain why adsorption as well as infection capacities were stronger for all RboP WTA-carrying *S. aureus* strains, while ΦK displayed lower adsorption and plating efficiency in *S. epidermidis* 1457. The myovirus ΦStab20 RBP1, on the other hand, only bound to *tarM*-deficient *S. aureus*. This difference is likely due to a difference in the amino acid sequence of the two RBP1 proteins, as shown in the binding assay (Figure 3a). Albeit present in the same cluster, the RBP1 of ΦK and ΦStab20 belong to different subclusters (Figure 1a, f & g). Across the different myovirus RBP subclusters we observed a variety of RBP binding patterns that seemed to correlate with the behavior of the respective phages (Figure 3a). For some myoviruses such as ΦPG-2021_10, the binding behavior of RBP2 did not match the phage infection pattern. Here, only binding to *S. aureus* Δ*tarM* Δ*tarS* with unglycosylated WTA could be observed, while RBP1 as well as the phage itself bound only to the *tarM*-deficient mutant.

Overall, the binding capacities of the various RBPs largely matched the binding and infection behavior of the full phages. To enable quick detection of phage RBPs in genomes of *S. aureus* phages and prediction of their WTA-dependent host range, we developed the **Ph**age **A**ureus **R**BP **I**dentification **S**ystem “PhARIS” detection tool, available via https://github.com/JKrusche1/PhARIS. This tool allows users to upload already known as well as uncharacterized *S. aureus* phage genomes, and predict their potential RBPs, the most similar RBP in the RBP cluster analysis, as well as its binding specificity based on similarity to already characterized RBPs.

### Φ13-RBP can be used to rapidly detect RboP WTA in a strain-specific manner

RboP WTA is found in almost every *S. aureus* and it can contribute to the pathogenicity of rare RboP-carrying NAS as recently demonstrated for *S. epidermidis* E73 (Du et al., 2021). Because of the important role of RboP WTA for virulence and horizontal gene transfer (Du et al., 2021), it is vital to detect RboP in other staphylococci. The RBP of Φ13, for which no second RBP was found, showed high affinity for GlcNAc-glycosylated RboP WTA, and moderate affinity for unglycosylated RboP WTA, while there was no affinity for GroP WTA (Figure 3).

Based on this finding, we used the RBP of Φ13 to assess the presence of RboP in many staphylococcal species (Figure 4a). Φ13-RBP binding was detected in close relatives of *S. aureus* such as *Staphylococcus schweitzeri* and *Staphylococcus argenteus*, as well as the more distantly related *Staphylococcus equorum*, *Staphylococcus arlettae*, *Staphylococcus saprophyticus*, and in most strains of *Staphylococcus xylosus,* suggesting that these strains produce RboP WTA. Strain-specific differences in Φ13-RBP binding could be detected for a variety of *Staphylococcus* species suggesting that they differ in the presence of RboP WTA. Most *S. epidermidis* carry GroP WTA decorated with glucose residues (Beck et al., 2024), but some clonal groups such as *S. epidermidis* ST10, ST23, and ST85 have recently been found to produce both, GroP and RboP WTA (Du et al., 2021). We could confirm this finding using the Φ13-RBP as it bound to *S. epidermidis* E73, while showing no adsorption to *S. epidermidis* E73 Δ*tarIJLM* or *S. epidermidis* 1457 (Figure 4b). For many strains of *S. xylosus* we found evidence for RboP WTA in the cell wall, but strain LTH 6232 did not bind to the Φ13-RBP (Figure 4b). Conversely, most of the tested *S. warneri* strains did not seem to carry RboP WTA, except for the clinical strain 197-70259724 (Figure 4b). Additionally, RboP WTA dependent Φ13-RBP binding was not observed in various other species including *Staphylococcus capitis, Staphylococcus hominis, Staphylococcus simulans*, *Staphylococcus pseudintermedius*, *Staphylococcus haemolyticus,* and *Bacillus subtilis* 168 (Figure 4a).

**Figure 4:**
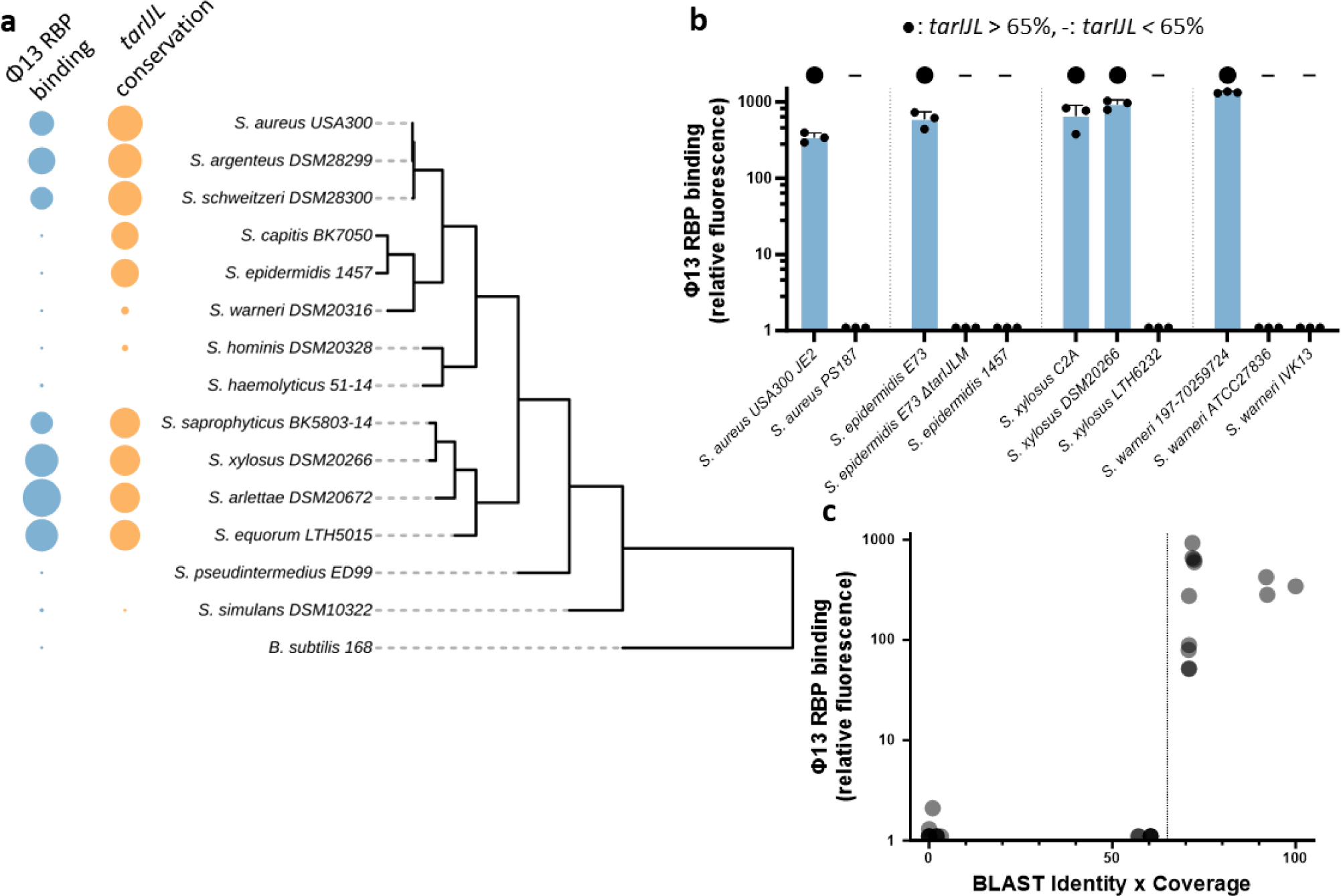
Φ13-RBP is a viable tool to detect RboP WTA in a wide range of staphylococci. **a,** Putative presence of RboP WTA as indicated by Φ13-RBP binding, and *tarIJL* cluster conservation (blast identity-x-coverage) for different *Staphylococcus* species and isolates. Phylogenetic tree based on 16s rDNA sequence. **b,** Fluorescence of different strains of *S. aureus, S. epidermidis, S. xylosus,* and *S. warneri* after co-incubation with GFP-coupled Φ13-RBP, after subtracting the background fluorescence of untreated bacteria. When lower than 1.1, the resulting value was set to 1.1 for visualization purposes. Presence of *tarIJL* with above 65% identity-x-coverage value was indicated with ●, absence with -. **c,** Fluorescence of GFP-coupled Φ13-RBP coincided with the presence of *tarIJL* in the genome of the corresponding bacteria. *S. aureus* USA300 JE2 *tarIJL* was used as query, and the resulting identity percentage was multiplied with the coverage percentage. Each dot represents one tested bacterial strain. All strains can be found in Table S6 and Figure S4. Strains with identity-x-coverage scores higher than 65% (marked by the horizontal line) were always bound by the Φ13-RBP, while strains with lower scores did not appear to have any RboP WTA. Full data can be found in Table S6.

These strain-specific differences prompted further investigation into the correlation between Φ13-RBP binding and presence of the *tarIJL* gene cluster. The *tarIJL* genes are necessary for the production of RboP WTA in *S. aureus* (Qian et al., 2006) and in *S. epidermidis* E73 (Du et al., 2021). BLAST analysis was used to identify *S. aureus* USA300 JE2 *tarIJL* homologous genes in the genomes of all tested strains. Similarity was quantified by multiplying BLAST identity percentage by the BLAST coverage percentage (Figure 4c). Many strains such as *S. epidermidis* 1457 or *S. capitis* carried parts of the *tarIJL* cluster in their genome but their *tarL* gene was substantially different, resulting in lower alignment coverage (around 80%). This difference coincided with the absence of Φ13-RBP binding and, presumably, RboP WTA. Indeed, the *tarIJL1* gene cluster of *S. epidermidis* 1457 has been found to be nonfunctional (Du et al., 2021). In total, RboP WTA could be detected at identity-x-coverage values above 65%, while genomes with lower *tarIJL* similarity were not able to bind Φ13-RBP (Figure 4c, b: ● for *tarIJL* > 65%, - for *tarIJL* < 65%). The strong correlation between Φ13-RBP binding and presence of a well-conserved *tarIJL* cluster suggests that it is possible to estimate if a specific *Staphylococcus* strain carries RboP WTA via evaluation of the identity-x-coverage score related to *S. aureus tarIJL*. Such a correlation could only be demonstrated for the genus *Staphylococcus* but not for other genera, as for example the RboP WTA carrying *Bacillus spizizenii* W23 (Brown et al., 2010), which was bound by the Φ13-RBP (Figure S4), carries a *tarIJL*-related gene cluster, while the identity-x-coverage score was only 34.6%.

## Discussion

*S. aureus* phages use RBPs to bind WTA and attach specifically to their host cells (Leprince & Mahillon, 2023), thereby targeting a vital pathogenicity factor, whose glycosylation pattern (“glycocode”) is known to differ among strains of *S. aureus* (Brown et al., 2013; van Dalen et al., 2020). This study elucidates the binding pattern of many different staphylococcal phages by assessing the binding of fluorescently labelled RBPs and correlating these to the phage infectivity and the presence of the RboP WTA biosynthesis encoding gene cluster *tarIJL* in the host-strain genomes. With the help of phylogenetic protein analysis, we show that many RBPs of *S. aureus* are closely related and can be separated into clusters and subclusters with distinct binding preferences (Figure 1). The high intra-cluster similarity may reflect the conserved architecture of the specific WTA glycotypes. While in unrelated species such as *E. coli,* phage RBPs continuously evolve as the host attempts to escape phage binding by mutating proteinaceous receptors such as LamB (Chatterjee & Rothenberg, 2012), glycosylated WTA in *S. aureus* appear to be very stable, as changes in WTA structure would require major changes in glycosyltransferase activity (Tamminga et al., 2022). Accordingly, *S. aureus* phage RBPs are probably not subjected to major selection pressure, which might explain the high similarity of RBPs of the individual clusters. Nonetheless, our data clearly indicate that TarM does not only protect *S. aureus* from podovirus infection (Li et al., 2015; Yang et al., 2023), but can also prevent infection by various myoviruses (Figure 3b). This role underlines the importance of TarM in the evolutionary defense against phages.

We also found that many *S. aureus* phages carry two different RBPs, one of which is probably responsible for initial reversible attachment while the other subsequently tethers the phage more stably to the host cells (Figure 1, Figure 2). Phages specific to other bacterial species often have more than one RBP to detect and bind their host. For example, the c2 phages of *Lactococcus lactis* first bind to carbohydrates on the cell surface and then to a membrane-attached protein (Tremblay et al., 2006). *S. aureus* phages, however, even when carrying two RBPs, only utilize WTA as binding epitope for both RBPs. As phages of other bacterial species, many *S. aureus* phages use one RBP to initially detect cells and approach the host in a reversible manner, while the second RBP mediates irreversible adsorption, after which infection can follow. The exact interplay of reversible and irreversible phage binding to WTA is yet unclear and necessitates further research.

The binding specificity of the Φ11 RBP1 in our study (Figure 1b, d) differed from that reported in previous work as this RBP bound to the Δ*tarM* Δ*tarS* mutant, while previous studies could not detect any binding of the protein to this strain (Li et al., 2016). While Li et al. used an RBP biotin labelling assay with high background fluorescence signal and WTA-glycosyltransferase mutants of strain RN4220, which produces lower amounts of WTA, we used a more sensitive assay and mutant strains in the USA300 JE2 background, which produces higher amounts of WTA (Wanner et al., 2017).

Based on the extreme heterogeneity in the sequence of RBPs from myoviruses that have been annotated to infect either *S. aureus* or NAS, we speculate that the RBPs of *S. aureus* and *S. epidermidis* myoviruses developed from a common ancestor, and that many myoviruses can infect both *S. aureus* and NAS, as previously shown (Goller et al., 2021). It is possible that small changes in RBP structure can lead to substantial changes in RBP binding specificity, as shown for myovirus ΦSA012, where single amino acid replacements in the RBPs can alter the infection behavior of the phage (Takeuchi et al., 2016). Such functional variations could explain the strong binding differences between myovirus RBP subclusters and should prompt further research into the variation of the RBP sequences (Figure 3b). The WTA binding site of these RBPs is yet unknown and could be elucidated by co-crystallization of the different RBPs with RboP WTA as it has been done for WTA-antibody and WTA-glycosyltransferase complexes (Di Carluccio et al., 2022; Gerlach et al., 2018).

Only myovirus RBPs were able to bind to both, *S. aureus* and *S. epidermidis*, while none of the *S. aureus* sipho- or podovirus RBPs was found to adsorb to *S. epidermidis* 1457 (Figure 3b). The lower adsorption and binding efficiency of ΦK to *S. epidermidis* 1457 compared to *S. aureus* possibly results from RBP2, as this protein had only very low or no affinity for *S. epidermidis* 1457 (Figure 3a). The fact that transducing siphoviruses can only bind either *S. aureus* or NAS but never both, has strong implications for horizontal gene transfer between NAS and *S. aureus*. It is still unclear how antibiotic resistance genes, such as *mecA*, have been transferred from GroP WTA carrying NAS such as *S. epidermidis* to RboP WTA carrying *S. aureus,* considering that phage transduction is regarded as the major way of horizontal gene transfer between these groups (Rolo et al., 2017). The results presented here show that nearly all known *S. aureus* podo- and siphoviruses can be grouped into one of four RBP1 clusters (Figure 1), and that the members of these clusters do not have the ability to bind to GroP WTA carrying strains such as *S. epidermidis* 1457 (Figure 3b). In contrast, all RBP1 proteins of myoviruses were able to adsorb to GroP WTA, and most were able to bind at least one *S. aureus* strain with RboP WTA. Thus, interspecies horizontal gene might be mediated by myoviruses. Additionally, giant viruses, which are unrelated to all other phage groups, seem to have an equally wide host range as many myoviruses and might be involved in gene transfer across species barriers (Uchiyama et al., 2014). However, until now, there has been no observation of transduction capacities for staphylococcal myoviruses. As transduction requires packaging of host DNA in the phage particles prior to bacterial lysis, we speculate that the activity of nucleases, highly enriched in the genomes of myoviruses, as well as the lytic nature of myoviruses may prevent promiscuous packaging and thereby interspecies horizontal gene transfer (O’Flaherty et al., 2004). Nonetheless, myoviruses carrying only few endonucleases could, in principle, transfer DNA between staphylococci. We cannot rule out the possibility that as-yet unknown NAS-infecting siphoviruses could additionally bind to *S. aureus* and thus transfer resistance genes, or that one of the non-classified *S. aureus*-infecting phages carry special RBPs with the ability to bind *S. aureus* and *S. epidermidis*.

Another avenue of interspecies horizontal gene transfer could be based on the existence of *Staphylococcus* strains that carry both GroP- and RboP WTA, such as *S. epidermidis* E73, and could therefore be infected by both *S. aureus* and NAS-specific phages. Some NAS strains other than E73 were bound by the RBPs of both, siphovirus Φ13 (RboP binding) and myovirus ΦBS1, which was found to bind only to some glycotypes of GroP WTA (Figure S4). These finding suggest that these NAS strains may express both RboP- and GroP WTA and can be infected by different phages, thereby functioning as a hub for horizontal gene transfer between staphylococci with different WTA backbone structures.

Combining Φ13-RBP binding assays with genome mining to detect the presence of the *tarIJL* cluster, we identified NAS species that are likely to carry RboP WTA (Figure 4a, c). In contrast to most *S. warneri* species, which did not carry *tarIJL*, the clinical isolate *S. warneri* 197-70259724 was bound by the Φ13-RBP and probably carried RboP WTA, suggesting that the pathogenicity of the opportunistic pathogen *S. warneri* might be shaped by changes in the WTA backbone structure (Figure 4b). Further investigations into the pathogenicity of rare RboP WTA carrying strains is necessary and could help understand the interaction between staphylococcal WTA and host receptors during infections. In this context, the Φ13-RBP could be a viable tool to find RboP WTA carrying species and strains even in distantly related Firmicutes such as *Bacillus spizizenii* or *Listeria monocytogenes* in a high-throughput fashion (Figure S4). Additionally, Φ13-RBP could be useful for optimization of drug delivery, as Φ13-RBP coupled antibacterial lysins might permit the specific targeting of RboP WTA carrying bacteria with high virulence potential (Zampara et al., 2020).

Overall, the host range of *S. aureus* phages was highly associated with RBP binding (Figure 3a), and the phages could be grouped into separate clusters based on the high RBP conservation. This association enables host range prediction of a given phage based on its RBP sequence by assignment to the closest intra-cluster RBP with known host range. To simplify such a phage host range analysis pipeline, the PhARIS toolbox was developed, which first identifies the RBP, followed by prediction of its binding capabilities, via genomic comparison. This strategy enables the investigation of the host range of newly isolated *S. aureus* phages, as well as some of the *S. epidermidis* myoviruses. Using this approach, we were able to find phages with similar host range as myovirus ΦStab20 and we could show that *agr-*mediated downregulation of *tarM* can impact infection of *S. aureus* by multiple different myoviruses (Yang et al., 2023). These myoviruses show only low overall genome similarity to ΦStab20 but were found to carry RBPs with high similarity to the ΦStab20 RBPs, and their infection behavior matched that of ΦStab20. The PhARIS tool can also be used to predict the strain specificity of phages and therefore help to design more specific and effective phage cocktails for phage therapy.

## Materials and Methods

### Bacterial strains and growth conditions

*Staphylococcus* and *Bacillus* strains were grown at 37°C on an orbital shaker in Tryptic Soy Broth (TSB). *E. coli* strains were grown in Lysogeny Broth (LB) at 37°C. *Staphylococcus* and *Bacillus* species were grown without antibiotics, while *E. coli* overexpression strains were grown with 10 µg/mL kanamycin. All strains, phages, and the respective propagation hosts can be found in Table S1.

### Phage propagation

Phages were propagated by inoculating a liquid culture to OD 0.4 of the phage-specific propagation strain (see Table S1) in 9 mL TSB + 5 mM CaCl_2_ and incubated at 37°C for 30 minutes. Next, 1 mL of phage lysate (titer 10^8^-10^10^) was added, and the mixture was incubated at 37°C for 4 hours (podo- and siphoviruses) or 30°C for 6 hours (myoviruses). Then, the lysate was centrifuged (5,000 x g for 5 minutes) and the supernatant was sterile filtered (0.45 µm) and stored at 4°C.

### Cloning of RBP-expressing *E. coli*

The His(6)-GFP-encoding DNA sequence was cloned in pET-28a(+) via Gibson assembly (Gibson et al., 2009), and used to transform chemically competent *E. coli* DC10B via heat shock for 30 seconds at 42°C. The RBP genes were either amplified by PCR from the phage genome or ordered as synthetic DNA fragments from Thermo Scientific via the Invitrogen GeneArt Synthesis Services in cases where the phage itself was unavailable as PCR template. Ordered primers and DNA fragments can be found in Table S4 & S5. The RBP-encoding fragments were then ligated onto the C-terminal end of the GFP-encoding fragment in the pET-28a(+)_GFP vector via Gibson assembly and used once again to transform *E. coli* DC10B. From there, resulting plasmids were purified and transferred by heat shock into chemically competent *E. coli* BL21(DE3) for protein overexpression.

### Cloning in *S. aureus*

Cloning of glycosyltransferase mutants in *S. aureus* USA300 JE2 was performed as previously described for RN4220 Δ*tarM* and RN4220 Δ*tarS* (Winstel et al., 2013). The marker-less deletion mutants of the glycosyltransferase genes *tarM* and *tarS*, as well as a Δ*tarM* Δ*tarS* double mutant, were originally generated by Gibson cloning of the two flanking regions of the respective genes in the mutagenesis vector pBASE6 or pKOR1. These original mutagenesis vectors were transferred by electroporation (pBASE6-tarM) and transduction via ϕ11 (pKOR1-TarS) into USA300 JE2, following the recombination procedure described elsewhere (Bae & Schneewind, 2006). The mutants were confirmed genotypically by PCR and controlled for *agr* activity and toxin production by cultivation on blood agar plates (Adhikari et al., 2007; Cheung et al., 2012), coagulase activity (using Biomerieux STAPH-ASE (Ref 55181)), and deposition of anti-WTA-Fabs (using clones 4461 and 4497 patent (Driguez et al., 2017)) as described previously (van Dalen et al., 2019). The strains, plasmids and oligonucleotides used are listed the supplemental information in Table S1, Table S3 and Table S4, respectively.

### Overexpression and purification of phage receptor-binding proteins

*E. coli* BL21(DE3) carrying the respective pET-28a(+)_GFP_RBP plasmids (See Table S2) were incubated overnight in LB at 37°C shaking. Overexpression cultures were inoculated to OD_600_ 0.1 in TSB and incubated for 2-3 hours at 37°C under shaking at 200 rpm, whereafter the temperature was shifted to 20°C for 15 minutes. Subsequently the overexpression was induced by addition of 1 µg/mL Isopropyl β-D-1-thiogalactopyranoside (IPTG) and incubated on a shakier overnight at 20°C.

The cells were collected by centrifugation (10 minutes at 4,000 x g), resuspended in lysis buffer (30 mM Tris-HCl pH 8.3 with 20 mM imidazole and 300 mM NaCl), treated with lysozyme, Triton X-100, and protease inhibitor tablets, and lysed by sonication. After removal of cell debris by centrifugation (17,000 x g, 10 min), the supernatant was sterile-filtered (0.22 µm), and the proteins were purified via nickel column chromatography. Smaller impurities and lysis buffer contents were removed by dialysis with Slide-A-Lyzer G3 dialysis cassettes (10K MWCO) Cat. Nr. A52971 overnight in RBP buffer (30 mM Tris-HCl pH 8.3 with 150 mM NaCl) at 4°C. Protein concentration was measured after dialysis via the Qubit Protein Assay.

### Spot assay

The phage count was enumerated as plaque forming units (PFU) per mL via agar overlay method (Kropinski et al., 2009). Briefly, *S. aureus* and *S. epidermidis* cultures were inoculated from overnight cultures at an OD_600_ of 0.05 in 4 mL of 0.5% TSA soft agar (TSB with 0.5% agarose). The soft agar was poured onto TSA plates. After solidification, 5 µL of dilution series of the phages were spotted onto each soft agar plate containing a different bacterial strain and incubated overnight at 37°C (sipho- and podoviruses) or 30°C (myoviruses).

### Adsorption assay

The phage adsorption assay was performed as described in (Xia et al., 2011) with slight modifications. Briefly, 100 µL of phage lysate containing 3*10^7^ or 3*10^8^ phages was mixed with 200 µL of bacteria (OD_600_ 0.5) and incubated for 10 minutes at 30°C with shaking (300 rpm). Afterwards, the bacteria and bound phages were removed by centrifugation at 13.000 x g for 5 minutes at 4°C and subsequent filtration of the supernatant (0.45 µM). The remaining phage lysates were serially diluted and then used in a spot assay for enumeration. The percentage of bound phages was calculated as ratio compared to the negative control that contained no bacteria during the initial incubation step.

### RBP flow cytometry assay

Overnight cultures of the test strains were washed once in RBP buffer (30 mM Tris-HCl pH 8.3 with 150 mM NaCl) and then diluted to OD_600_ 0.4. 30 µL of bacteria were mixed with 30 µL of the purified RBPs (0.2 µM) and incubated for 8 minutes at 20°C shaking (350 rpm). Next, the bacteria were washed by addition of 90 µL RBP buffer followed by centrifugation for 2.5 min at 8,000 x g. The bacteria were resuspended in 150 µL fresh RBP buffer. The bacterial suspension was then transferred into FACS tubes and GFP-mediated fluorescence of the cells was measured via flow cytometry in a BD FACSCalibur (FL1).

### RBP fluoreader assay

Overnight cultures of the test strains were washed once in RBP buffer (30 mM Tris-HCl pH 8.3 with 150 mM NaCl) and then diluted to OD_600_ = 2. 30 µL of bacteria were mixed with 30 µL of the purified RBPs (2 µM) and incubated for 4 minutes at 20°C with shaking (350 rpm). Next, the bacteria were washed by centrifugation for 2.5 min at 8,000 x g, whereafter the bacteria were resuspended in 150 µL RBP buffer. The fluorescence of the samples was measured in a 96-well plate in a BMG CLARIOstar (Ex: 470-15; Em: 515-20).

### Fluorescence microscopy

Bacteria and phage RBPs were processed as in the RBP FACS assay. After washing, 32 µL of the bacterial suspension was transferred into µ-Slide 15 Well 3D (formerly µ-Slide Angiogenesis) ibidi Cat.No:81506 and centrifuged twice for 6 minutes at 600 x g. The supernatant was carefully removed, and the wells were filled with 10 µL ibidi mounting medium Cat.No:50001. The samples were analyzed using a fluorescence microscope.

### RBP clustering

A curated RBP list was created by *in-silico* analysis of the genomes of ΦP68, ΦCSA13, Φ11, Φ12, Φ13, ΦK and ΦRemus. These phages were chosen as representatives of their respective cluster due to availability of published data or highest RBP similarity to other phage RBP clusters. The staphylococcal phage database was created by isolation of all phages in the INPHARED database (Cook et al., 2021) that contained the keywords “staph”, “aureus” or “epidermidis”. The curated RBPs were then protein-BLASTed in the staphylococcal phage database, and results were filtered with a cutoff of at least 65% overlap and 65% identity. The results were used for ClustalOmega MSA, which was then processed in SimplePhylogeny and visualized in iTOL (Letunic & Bork, 2021; Sievers et al., 2011). The interactive RBP1 tree is accessible under https://itol.embl.de/tree/4652552332411677924980. The interactive RBP2 tree is accessible under https://itol.embl.de/tree/46525523312761688382866.

### In-silico methods

Clustering was performed with protein blast (Altschul et al., 1997), ClustalOmega (Sievers et al., 2011), SimplePhylogeny (Madeira et al., 2022), and interactive tree of life (iTOL) (Letunic & Bork, 2021). Analysis of FACS data was done with FlowJo 10.0. Visualization of data and statistical analysis was performed with GraphPad Prims 10.0. PhARIS was developed with Spyder 5.5.0 in Python 3.12.0, the source code can be accessed on GitHub https://github.com/JKrusche1/PhARIS.

## Acknowledgements

We thank M. Skurnik and H. Ingmer for providing phage Stab20, M. Lössner for providing phage 3A, *L. monocytogenes* EGDe and *L. monocytogenes* EGDe Δ*rmlB*, E. Gómes-Sanz for providing phages PG-2021_10, PG-2021_17 and their respective host strains, and F. Götz for providing *S. xylosus* LTH6232. We thank L. Lo Presti for editorial assistance and N. Vetter for assistance with fluorescence microscopy. A.P. acknowledges financial support from Deutsche Forschungsgemeinschaft, (SPP 2330 and PE 805/7-1) and infrastructural funding from the Cluster of Excellence EXC 2124 ‘‘Controlling Microbes to Fight Infections’’ project ID 390838134.

## Author contributions

Conceptualization, J.K., C.B. and A.P.; methodology, J.K., C.B., E.L, D.G., C.W. and A.P.; investigation, J.K. and E.L.; formal analysis, J.K.; visualization, J.K.; writing–original draft, J.K.; writing–review & editing, J.K., C.B, E.L., D.G., and A.P.; supervision, A.P., funding acquisition, A.P.

