## Supplementary information for "Systematic classification of phage receptor-binding proteins predicts surface glycopolymer structure in *Staphylococcus* pathogens"

**Supplement**

**Table S1:** Bacteria used in this study

| **Bacterial strain** | **Description** | **Source** |
|---|---|---|
| *S. aureus* 8325-4 ϕ13-kana | Phage propagation | (Tang et al., 2017) |
| *S. aureus* RN4220 | Phage propagation & host range determination | (Kreiswirth et al., 1983) |
| *S. aureus* RN4220 Δ*tagO* | Host range determination | (Xia et al., 2011) |
| *S. aureus* RN4220 Δ*tarM* | Phage propagation & host range determination | (Winstel et al., 2013) |
| *S. aureus* RN4220 Δ*tarS* | Host range determination | (Winstel et al., 2013) |
| *S. aureus* RN4220 Δ*tarM* Δ*tarS* | Host range determination | (Winstel et al., 2013) |
| *S. xylosus* M1997-2/10 | Phage propagation | (Goller et al., 2021), Obtained from Martin Lössner, Zurich |
| *E. coli* BL21(DE3) | RBP overexpression (See Table S3) | New England Biolabs |
| *S. aureus* USA300 JE2 WT | RBP specifity determination | NARSA strain collection |
| *S. aureus* USA300 JE2 Δ*tagO* | RBP specifity determination | (Slavetinsky et al., 2023) |
| *S. aureus* USA300 JE2 Δ*tarM* | RBP specifity determination | This study |
| *S. aureus* USA300 JE2 Δ*tarM* Δ*tarS* | RBP specifity determination | This study |
| *S. aureus* USA300 JE2 Δ*tarS* | RBP specifity determination | This study |
| *S. epidermidis* 1457 | RBP specifity determination | (Mack et al., 1992) |
| *B. subtilis* 168 | Φ13-RBP RboP testing | DSMZ |
| *B. spizizenii* W23 | Φ13-RBP RboP testing | DSMZ |
| *L. grayi* ATCC25401 | Φ13-RBP RboP testing | |
| *L. monocytogenes* EGDe | Φ13-RBP RboP testing | (Glaser et al., 2001), Obtained from Martin Lössner, Zurich |
| *L. monocytogenes* EGDe Δ*rmlB* | Φ13-RBP RboP testing | (Eugster et al., 2015), Obtained from Martin Lössner, Zurich |
| *S. argenteus* DSM28299 | Φ13-RBP RboP testing | DSMZ |
| *S. arlettae* DSM20672 | Φ13-RBP RboP testing | DSMZ |
| *S. aureus* PS187 | Φ13-RBP RboP testing | (Winstel et al., 2014) |
| *S. aureus* PS187 Δ*tagN* | Φ13-RBP RboP testing | (Winstel et al., 2014) |
| *S. capitis* BK7050 | Φ13-RBP RboP testing | This study |
| *S. epidermidis 1457* Δ*tagE* | Φ13-RBP RboP testing | (Beck et al., 2024) |
| *S. epidermidis* E73 | Φ13-RBP RboP testing | (Du et al., 2021) |
| *S. epidermidis* E73 Δ*tarIJLM* | Φ13-RBP RboP testing | (Du et al., 2021) |
| *S. equorum* LTH5015 | Φ13-RBP RboP testing | (Li et al., 2015) |
| *S. haemolyticus* 51-14 | Φ13-RBP RboP testing | (Winstel et al., 2015) |
| *S. hominis* DSM20328 | Φ13-RBP RboP testing | DSMZ |
| *S. pettenkoferi* 210-70229632 | Φ13-RBP RboP testing | This study |
| *S. pseudintermedius* ED99 | Φ13-RBP RboP testing | (Bannoehr et al., 2007) |
| *S. saprophyticus* 47/1 | Φ13-RBP RboP testing | This study |
| *S. saprophyticus* 47/2 | Φ13-RBP RboP testing | This study |
| *S. saprophyticus* 78-70165600 | Φ13-RBP RboP testing | This study |
| *S. saprophyticus* BK5803-14 | Φ13-RBP RboP testing | (Winstel et al., 2015) |
| *S. saprophyticus* BK6292/13 | Φ13-RBP RboP testing | (Li et al., 2015) |
| *S. saprophyticus* VA213751/14 | Φ13-RBP RboP testing | This study |
| *S. schweitzeri* DSM28300 | Φ13-RBP RboP testing | DSMZ |
| *S. simulans* ATCC27848 | Φ13-RBP RboP testing | DSMZ |
| *S. simulans* NT219 | Φ13-RBP RboP testing | (Cramton et al., 1999) |

| | | |
|---|---|---|
| *S. warneri* 197-70259724 | Φ13-RBP RboP testing | This study |
| *S. warneri* ATCC27836 | Φ13-RBP RboP testing | DSMZ |
| *S. warneri* BK15472/12 | Φ13-RBP RboP testing | (Winstel et al., 2015) |
| *S. warneri* BK6091/14 | Φ13-RBP RboP testing | This study |
| *S. warneri* BK9351/13 | Φ13-RBP RboP testing | This study |
| *S. warneri* IVK13 | Φ13-RBP RboP testing | This study |
| *S. warneri* IVK51 | Φ13-RBP RboP testing | This study |
| *S. warneri* VA16066/14 | Φ13-RBP RboP testing | This study |
| *S. xylosus* C2A | Φ13-RBP RboP testing | (Gotz et al., 1983) |
| *S. xylosus* DSM20266 | Φ13-RBP RboP testing | DSMZ |
| *S. xylosus* LTH6232 | Φ13-RBP RboP testing | Obtained from Friedrich Götz, Tübingen |

**Table S2:** Phages used in this study

| **Phage** | **Propagation/induction strain** | **Source** |
|---|---|---|
| ΦP68 | *S. aureus* RN4220 Δ*tarM* | (Li et al., 2015) |
| Φ11 | *S. aureus* RN4220 Δ*tarM* | (Novick, 1967) |
| Φ80α | *S. aureus* RN4220 Δ*tarM* | (Novick, 1967) |
| Φ3A | *S. aureus* RN4220 Δ*tarM* | Obtained from Martin Lössner, Zurich |
| Φ47 | *S. aureus* RN4220 | (Ralston & Baer, 1964) |
| Φ13 | 8325-4 ϕ13-kana | (Tang et al., 2017) |
| ΦN315 | *S. aureus* RN4220 | (Gerlach et al., 2018) |
| ΦK | *S. aureus* RN4220 | (O'Flaherty et al., 2005) |
| ΦStab20 | *S. aureus* RN4220 Δ*tarM* | (Oduor et al., 2019), Obtained from Hanne Ingmer, Copenhagen |
| ΦPG-2021_10 | *S. xylosus* M1997-2/10 | (Goller et al., 2021), Obtained from Martin Lössner, Zurich |
| ΦPG-2021_17 | *S. xylosus* M1997-2/10 | (Goller et al., 2021), Obtained from Martin Lössner, Zurich |

**Table S3:** Plasmids used for RBP overexpression

| **Plasmid** | **Insertion** | **Progenitor plasmid** |
|---|---|---|
| pET-28a_GFP | His(6)-SSG-eGFP-GSGSGS inserted for RBP fusion | pET-28a(+) |
| pET-28a_CSA13-RBP | C-terminal ΦCSA13 RBP (151-647) | pET-28a_GFP |
| pET-28a_P68-RBP | C-terminal ΦP68 RBP (116-642) | pET-28a_GFP |
| pET-28a_11-RBP1 | C-terminal Φ11 RBP1 | pET-28a_GFP |
| pET-28a_11-RBP2 | C-terminal Φ11 RBP2 (241-607) | pET-28a_GFP |
| pET-28a_12-RBP1 | C-terminal Φ12 RBP1 (118-636) | pET-28a_GFP |
| pET-28a_12-RBP2 | C-terminal Φ12 RBP2 (248-499) | pET-28a_GFP |
| pET-28a_13-RBP | C-terminal Φ13-RBP (825-1225) | pET-28a_GFP |
| pET-28a_K-RBP1 | C-terminal ΦK RBP1 (50-458) | pET-28a_GFP |
| pET-28a_K-RBP2 | C-terminal ΦK RBP2 | pET-28a_GFP |
| pET-28a_Stab20-RBP1 | C-terminal ΦStab20 RBP1 (50-458) | pET-28a_GFP |
| pET-28a_SA012-RBP1 | C-terminal ΦStab20 RBP1 (50-458) | pET-28a_GFP |
| pET-28a_PG-2021_10-RBP1 | C-terminal ΦPG-2021_10 RBP1 (50-458) | pET-28a_GFP |
| pET-28a_PG-2021_10-RBP2 | C-terminal ΦPG-2021_10 RBP2 | pET-28a_GFP |
| pET-28a_PG-2021_17-RBP1 | C-terminal ΦPG-2021_17 RBP1 (50-458) | pET-28a_GFP |
| pET-28a_BS1-RBP1 | C-terminal ΦBS1 RBP1 (62-465) | pET-28a_GFP |
| pBASE6 tarM | cloning Δ*tarM* | pBASE6 |
| pKOR-tarY | cloning Δ*tarS* | pKOR |

**Table S4:** Primers used in this study

| Cloning primers | Sequence (5’-3’) |
|---|---|
| pET28a_fw | ACAAAGGTAGTGGTAGTGGTAGTTAAGATCCGGCTGCTAACAAAGC |
| pET28a_rv | TTTGCTAGCGCTCATGCCGCTGCTGTGATGATGATGATGATGG |
| GFP_fw | CATCACAGCAGCGGCATGAGCGCTAGCAAAGGAGAAG |
| GFP_rv | GATCTTAACTACCACTACCACTACCTTTGTATAGTTCATCCATGCCATGTGTAAT |
| pET28a_GFP_vector_fw | GATCCGGCTGCTAACAAAG |
| pET28a_GFP_vector_rv | ACTACCACTACCACTACCTT |
| 11_primary_fw | GAACTATACAAAGGTAGTGGTAGTGGTAGTATGAGTAATAAACTAATTACAGATTTAAGTAGAGTCTTTGACTACAG |
| 11_primary_rv | TTCCTTTCGGGCTTTGTTAGCAGCCGGATCTTATTCAACCACCTTTCCTTCGAATAAACTCC |
| 11_secondary_fw | TAGTGGTAGTGACCTTGTTAAAGGTAATTCAAC |
| 11_secondary_rv | TAACAAGGTCACTACCACTACCACTACC |
| 12_primary_fw | AAGGTAGTGGTAGTGGTAGTAATAAAGAGGAAACTATAAGAGAATTAAATAAGACCA |
| 12_primary_rv | GCTTTGTTAGCAGCCGGATCTTAACTTATAATTCTCCCTTCGTGTAAAG |
| 12_secondary_fw | TAGTGGTAGTGGTTTAGTCACAAGTGG |
| 12_secondary_rv | TGACTAAACCACTACCACTACCACTACC |
| 13_RBP_fw | GAACTATACAAAGGTAGTGGTAGTGGTAGTAGAGAGGGTCTTGATATCAATGT |
| 13_RBP_rv | TTCCTTTCGGGCTTTGTTAGCAGCCGGATCTTATCCTGCATTCTTTGACTCC |
| K_primary_fw | ACAAAGGTAGTGGTAGTGGTAGTAAAATTAAACTAACTGATGATAAAGGATTAACTAAAT |
| K_primary_rv | TTCGGGCTTTGTTAGCAGCCGGATCCTATGGCATATTAATACCTATAATTCTTGTAAC |
| K_secondary_fw | ATACAAAGGTAGTGGTAGTGGTAGTATGGCATTTAACTACACGC |
| K_secondary_rv | TTCGGGCTTTGTTAGCAGCCGGATCTTATCCTCTATTAATTCCCATAATATTGTATACCT |
| P68_RBP_fw | AAGGTAGTGGTAGTGGTAGTTTAACATCGGCATTTGCTTTAT |
| P68_RBP_rv | GCTTTGTTAGCAGCCGGATCCTATTTTTGATGTTTTGCTACC |
| PG-2021_10_primary_fw | TACAAAGGTAGTGGTAGTGGTAGTAAAATTAAACTAACTGATGATAAAGGATTAACTAAA |
| PG-2021_10_primary_rv | TTCGGGCTTTGTTAGCAGCCGGATCCTATGGCATATTAATACCTATAATTCGTG |
| PG-2021_10_secondary_fw | ATACAAAGGTAGTGGTAGTGGTAGTATGGCATTTAACTACACGC |
| PG-2021_10_secondary_rv | TTCGGGCTTTGTTAGCAGCCGGATCCTAAAGTGTGTTAATTCCCGC |
| Stab20_secondary_fw | ATACAAAGGTAGTGGTAGTGGTAGTATGGCATTTAACTACACGC |
| Stab20_secondary_rv | TTCGGGCTTTGTTAGCAGCCGGATCCTAAAGTGTGTTAATTCCTGC |
| TarMinsiderev | CAATTCGCTTCGTTGGTACCATTC |
| TarMoutsidefw | ATATTTAAAAATTATGAGCAGATAGAGCCA |
| TarMoutsiderev | ATCTTCACACTTAAACTTACAAGAGGATATG |
| TarSinsidefw | ATTTGATAATCATAAAATTGTTACAGAGCTAA |
| TarSoutsidefw | TGTTCCAGGTAAAATTGTGCAATC |
| TarSoutsiderev | GCTTTCGGATAATCAGTTACTACATCT |
| **Sequencing primers** | **Sequence** |
| spacer-GSGS | AGGTAGTGGTAGTGGTAGT |
| T7 | TAATACGACTCACTATAGGG |
| T7-term | CTAGTTATTGCTCAGCGGT |

**Table S5:** DNA fragments used for cloning of RBPs

| **Synthetic DNA fragments** |
| --- |
| >PG-2021_17_primary_RBP<br>ATACAAAGGTAGTGGTAGTGGTAGTAAAGCCTATGGAAATATAACTACTCTTAAGGACATAAAAG<br>AGCCAGGTTACTATTATATTAATGCAAGAGCCTTTGCAATGTTAACAGATAAACCTGATATAGAA<br>TCTATTGATGTAGTACTTCAGGTACTACCTTTAGAATCTTCTAACCGAGTAGTACAACACTTATA<br>TACTTTGTCTACTAATAATAGTCAAATTAAGACTATATATAGATATGTTTCAGGAAATTCAAGTT<br>CAGAGTGGCAGTTTATTCAAGGACTACCAAGTAATAAAAATTCTGTTATATCAGGTTCTAACTTA<br>GAAGATTTAACTTCCCCTGGGGTATACTTTGTAACAGGTATGACAGACGGTATGCCTGATGGAGT<br>TAGCTCAGGGTTCTTAGAATTAAATATAGATGCTAATGATAATAGAATAGCTAAGGTAACTGATA<br>TTGAAACAGGAAAAGAGTATACTAAAGTTAAGAAACCTACAGGGGTATACGCAGAATGGAAAAAA<br>GAGCTAGAACCTACGGATATGCAAAAGTATTTATTAAGTAGTATAAAGGATGATGGCAGTGTTAC<br>ATTTCCATTAATGGTTTACACTTCAAGTAATAAAACATTCCAACAAGCTGTTTTAGACCATGTTG<br>ATAGTACAGGGCAAACTACTTTTACATTCTATGTTCAAGGAGGGGTTCCAGGTTCACCTATGCCT<br>AACAGTTGCAGAGGTATTTTTATTTCAGATACATCAAATATTGCTAGTTTTCATGGTGTATACAC<br>TGCAGTAGGTACTGATGGTAGGGATGTTACAGGTTCAGTAGTAGGGGGTAATTGGACTACCCCTA<br>AAGCTTCTCCTTCCTATAAAGAATTATGGACTGGGGCACAATCATTTTCCTCTATTGGAACTTCC<br>AAGAAGTTAGAAGACGATATAAGTAATTACTCATACGTAGAGGTTTATACTAAACATAAAACTGT<br>AGAGAAAACTAAGGGAAATGATGACACAGGAACTATTTGCCATAAGTTTTATTTAGATGGCAGTA<br>AAACTTATGTTTGTTCAGGAACTTTTGTTTCAGGAGAAAAAACAGACAAAACAGTACCTGTTACA<br>GAATTTTATAGAGTAGGTATTAATTTTTCAGGAACAACTTGGAAAGTAGTAGATAGTGCAGTACA<br>AAATAATAAAACTCAATACGTCACAAGAATTATAGGTATTAATATGCCATAGGATCCGGCTGCTA<br>ACAAAGCCCGAA |
| > BS1_primary_RBP<br>GAACTATACAAAGGTAGTGGTAGTGGTAGTAACAAGACAATAAGGTACTTATCAGACTTAACTCA<br>AGCAGGTACTTACTATGTAACAGCTTCTGTACTAGCGACATTACCAGACAAACCTAAATCTCTAA<br>TTACAAGTGATTCTATCGTAGAAATAAAACCTACAAGAAGAGTAGATGAAGTTATTCAACAAATC<br>ACAACATTAGGTAGAGACTTAGAAGATGGTAAAGTTGTTTATAGGTTTATCTCATCAGAAGGTAA<br>CTCAGAATGGGTTTACTTTAATGCGTTATCTAATAACAGATACTTTAGTAAGAGTTTAGATTTAT<br>CAAATACAACTGAACCTGGAACTTATTTCTTAACAAATAATAGTACTGGGCTACCTGAAGAGATT<br>AAAAATGCTGAAGGTATTGTTAAAATATTTGTAGATAACAATAATAACAGATATTATGAATATTA<br>CAACAATGAATTAGGAATATTTTATACAGGGTATAAAACTTCGTCTCAGTCATCAATAGATTGGA<br>AGACGAATAAAACAGATGCAAGTGAAATAAAAAATTTCTTGATAGGAGATATGAGTGTTACAGAT<br>ACAGGTATAGAATATCCTACTTTCGGGTTGCTTACGTACACATCAGATAATAAAGATTGGGAGAC<br>AGCAGTAAAAGAAAAACTAGCTACAGGTAAGACAATGTTTACCTTTTATTGTCAAGGTGGTGTAG<br>ATAAATCACCCGCTGGTAAGTTTAGTAGTCGAGGAATGGTAATTTCTGATGTACCAGGAGGTAGT<br>TATGGTGTTTACTATGCAATTACTAATGGTGGTAAACTATTTACAGGAGCTATAAGTAATTATAC<br>TTTTACTACCCCAAACGTTACGCCAAATGGAAATATCTTATGGCAAGGTGCTTTAAACTTTAAAA<br>CTGTTGATAAAACTCAAAAAATGAGAGATACAATTAATAACTATGATTATGTTGAAATATATACT<br>AAAATGAGAACATATAAAGAAGCTAAAGGAAAAGACCTTATTCAAAATATAGCACATAAATTCTA<br>TAGAGATGGGGAAACTTATTTTGTATGTACAGGCTCTTTACTAGGTGGAGAGTTTGATGGATCTA<br>CTGCCCCTTCTCAAGATGTATATAGGGTAAGTTTAGAGTTTAAAGGAGATACTTTCCAACTTAAA<br>GATTCTGCAATCAATAATAGTAAAACTCAATATGTAACAAGGATTATTGGTTACAACATGTTAGA<br>TGCCTTTTAAGATCCGGCTGCTAACAAAGCCCGAAAGGAA |
| > CSA13_primary_RBP<br>GAACTATACAAAGGTAGTGGTAGTGGTAGTGGGTTCGAATTAAATGAGCTTGAACCTAAGTTTGT<br>AATGGGCTTTGGTGGTGTTCGTAACGCTGTAAACCAAAGTATATTAATTGATAGAGAAACAAATC<br>AAATGTACTCAACACAATCCGACTCACAAAAAAAAGAGGGTTTTTGGATTAACAAATTAACACCT<br>AGTGGTGATTTAATTTCTAGTATGCATATTGTAGAGGGTGGTCATGGTACTAATATTGCTTTAGA<br>AAGAAAAAGTAACGGTGAGATAAAAATATGGTTGTATCATGTAGGACTTTCAAAACTTGTACAAA<br>TCGCTTATAGAGATAATACAACATTAACAGTAGATGATGCAAAAGCATTAACTGATTTCACGCCA<br>GCATCTTCACACGAATATTTTACGATTATTATGGATGAAGAAAATGACAAATTAGTATTTAGATA |

TGGCAGTGGTTTAATACAAGTACGTTCAAGAGAAGATGTAATTAATCACATTGACAATGTAGAAA
AAGAACTACAAATTGATGTAAAAGAAAACACACCAGACAGACCTATGCAAGGTATTGCAGTTTAT
GGTGATGATTTATACTGGTTAAGTGGTAGGTCAGATTTAGATAGTAAAACATTAATTCAAAAATA
TAGTTTTAAAACAAAATCAAAAGTATACGATTATTATTTAAACGATGTTTCATTTGAACATGGTG
TTGAAAATCCTAGAGATAATTTTAGAGAGCCTGAGGGTATATATTTATACGTTAATCCTAGAACT
AAAAAACAATCTTTATTAATTGCATATACAACTGCTGGTGGTGGAAAAAGACAAAACTTATTGTA
TGGTTTCTTTCAAACTGGTGAATATGAACGTTTTGTTGCATTAACGTTAAAAGGTGGGCAAAACT
ATAAACTAACGAAAGATGACGGTAGAGCTTTATCAATAAGAGATGGTGTGACAAGTTTAAGCCAA
ATTACCGAAACTGGATTTTATTATATTATGACAGAACATACTAAAGTTATGGATGATTTTCCATA
TATCCATGATGATGCAGGCTGGTTTTTATTTGTATCACCAAAAACACAACAGTTAGGTGGATATC
AAGAATTAACAAGAAATTCAGGATTAAGAAAAATGCTAAAACTAATAAGAAGTTTCACATATAAC
TTAGATAAACAAAAATTTGAATTTGGTAGCTGGTCGGTTATTGATACTAATTCAACTGAAAAAGA
ATACGTTGAGGCTAAATATTTTGATTATTGGATTGCTAAAATTACATTACCTGGAGAATATTACA
TTACTTCAACACAAATGGACCAATTTAAAGATAAACCAGGGGGTATTGGCGAAGCTGGTGCGTGG
TTACAAGTATCAAGTGGTAACGTTTCAGGTGAAGTAAGACAAACGTTATTAATAAATTCATCAAC
ATATAAAGAGTTTTATAGTCTTGTAAGAGTAAAAAATACACATCGTGAGTATGATTGGGTAGCAA
AACATCAAAAATAGGATCCGGCTGCTAACAAAGCCCGAAAGGAA

>SA012_primary_RBP
GAACTATACAAAGGTAGTGGTAGTGGTAGTAAAGTTAAATTAACAGATGATAAGGGTTTATCTAA
ATCTTATGGGAATATAACTGTAATTAGGGATATAAAAGAACCAGGTTATTATTATATAAATGCAA
GAACATTAGCTACATTATTAGATAAACCTGATATGGAATCAATAGATGTTTTACTCCATGTACTA
CCTTTAGATTCATCTAATAGAGTAATACAGCATATATATACATTGTCTACTAATAACAATCAAAT
TAAGACATTATATAGGTTTGTTTCGGGAAACTCTAGTTCAGAATGGCAGTTTATAACTGGATTAC
CTAGTAATAAAAATGCTGTTATTTCAGGAACTAATATCCTGGATATATCTTCACCAGGTATTTAC
TTTGTTATGGGAATGACAGGAGGAATGCCAAAAGGTGTAAGTTCGGGATTTTTAGATTTGAATAT
AGATGCTAATGATAATAGATTAGCTAAGTTAACTGATTCTGAAACAGGTAAAGAATACACTAGTA
TTAAAAAACCTACAGGTGTATACACAGAGTGGAAAAAAGAATTAGAACCTCAAGATATGCAAAAA
TATTTATTAAGTAGTATCAGGGATGATGGTAGTGCATCTTTCCCATTACTTGTGTACACTAGTGA
TAATAAGACTTTTCAACAGGCTGTTATAGACCATATAGATAGAACAGGGCAAACAACCTTTACCT
TTTATGTTCAAGGTGGTGTGACAGGTTCTCCTATGTCTAATAGCTGTAGAGGATTATTCATGTCA
GATACACCTAATACATCTAGTTTACATGGTGTTTATAATGCAGTAGGTACTGATGGTAGAAATGT
AACAGGCTCAGTAGTAGGTAGTAGTTGGACTTTTCCTAAAACATCACCGTCACATAAAGAATTAT
GGACGGGAGCACAATCATTCTTATCCACAGGTACTACTAAGAATTTAGCAGATGATATTAGTAAT
TATTCTTATGTAGAAGTTTATACTAAACATAAGACATCAGAGAAAACTAAAGGTAATGATGATAC
AGGTACAATTTGTCATAAATTTTATCTAGACGGTAGTGGTACCTACGTTTGCTCAGGTACATTTG
TATCAGGAGATAGAACCGATACAAAACCACCAATTACAGAATTTTATAGAGTAGGTGTATCCTTT
AAAGGTTCAACATGGACTCTTGTAGATAGCGCAGTGCAAAATAGTAAAAATCAATATGTTACAAG
AATTATAGGTATTAATATGCCATAGGATCCGGCTGCTAACAAAGCCCGAAAGGAA

**>**Stab20_primary_RBP
GAACTATACAAAGGTAGTGGTAGTGGTAGTAAAATTAAACTAACTGATGATAAAGGATTAACTAA
ATCTTATGGAAGCATAACAGCTCTTAGAGATATAAAAGAACCTGGTTACTATTATATTGGAGCTA
GAACATTAGCAACGTTATTAGATAAGCCTGACATGGAGTCTATTGATGTTGTATTACATGTAGTA
CCTCTTGACACTTCTAGTAAAGTAGTTCAACATTTATATACACTATCTACTAATAATAACCAAAT
TAAAATGTTATATAGATTTGTCTCAGGAAACTCTAGTTCAGAATGGCAATTTATTCAAGGATTAC
CTAGTAATAAAAATGCTGTTATATCAGGTACTAATATTTTAGATATATCTTCACCAGGTGTTTAT
TTTGTTATGGGAATGACAGGAGGTATGCCTAGTGGTGTAGACTCAGGTTTTTTAGATTTAAGTGT
AGATGCTAATGACAATAGACTAGCTAGGTTAACTGACGCTGAAACTGGTAAAGAGTATACTAGTA
TTAAGAAACCTACAGGAGTATACACAGCCTGGAAAAAAGAATTTGAACCAAAAGATATGGAGAAA
TATTTATTAAGTAGTATTAGAGACGATGGTAGTGCATCATTTCCACTTCTAGTTTACACTAGTGA
TAATAAAACATTCCAACAGGCTATTATAGACCATATAGATAGAACAGGTCAAACAACCTTTACTT
TCTACGTTCAAGGCGGTGTGTCAGGTTCACCTATGTCTAATAGTTGTCGAGGATTATTCATGTCT

GATACACCTAACACATCTAGTTTACATGGTGTCTATAATGCTATAGGTACGGATGGTAGAAACGT
AACAGGTTCAGTGGTAGGAGGTAACTGGACTTCACCAAAAACATCTCCTTCTCATAAAGAACTAT
GGACGGGAGCACAGTCATTCTTATCTACAGGGACTACTAAGAATTTATCAGATGATATCAGTAAC
TATTCTTATGTAGAGGTCTATACTAAACATAAGACAACAGAGAAGACTAAAGGTAATGATGATAC
AGGTACAATTTGCCATAAGTTTTATTTGGATGGCAGTGGTACTTATGTTTGTTCAGGTACATTTG
TATCGGGAGATAGAACAGATACAAAACCACCTATTACAGAGTTTTACAGAGTAGGTGTATCCTTC
AAAGGTTCAACATGGACTCTTGTAGATAGTGCAGTACAAAATAGTAAAACTCAATACGTTACAAG
AATTATAGGTATTAATATGCCATAGGATCCGGCTGCTAACAAAGCCCGAAAGGAA

**Table S6:** Results of discontinuous Megablast of NAS to *tarIJL* of S. aureus and Φ13-RBP binding [Median fluorescence]

| Strain | *tarIJL* identity | *tarIJL* coverage | ident. x cover. | Φ13-RBP binding |
|---|---|---|---|---|
| *B. subtilis* 168 | 0% | 0% | 0% | 1.1 |
| *S. pseudintermedius* ED99 | 0% | 0% | 0% | 1.1 |
| *S. warneri* BK6091/14 | 0% | 0% | 0% | 1.1 |
| *S. warneri* IVK13 | 0% | 0% | 0% | 1.1 |
| *S. simulans* DSM20322 | 96% | 1% | 1% | 2.1 |
| *S. hominis* DSM20328 | 93% | 2% | 2% | 1.1 |
| *S. aureus* PS187 | 71% | 3% | 2% | 1.1 |
| *S. aureus* PS187 ΔtagN | 71% | 3% | 2% | 1.1 |
| *S. warneri* DSM20316 | 80% | 4% | 3% | 1.1 |
| *S. capitis* BK7050 | 71% | 80% | 57% | 1.1 |
| *S. epidermidis* E73 ΔIJLM2 | 69% | 83% | 57% | 1.1 |
| *S. epidermidis* 1457 | 68% | 89% | 60% | 1.1 |
| *S. epidermidis* 1457 ΔtagE | 68% | 89% | 60% | 1.1 |
| *S. saprophyticus* VA213751/14 | 71% | 100% | 71% | 79.7 |
| *S. saprophyticus* 47/1 | 71% | 100% | 71% | 52.2 |
| *S. saprophyticus* 47/2 | 71% | 100% | 71% | 51.4 |
| *S. saprophyticus* 78-70165600 | 71% | 100% | 71% | 89.1 |
| *S. saprophyticus* BK5803-14 | 71% | 100% | 71% | 275.1 |
| *S. xylosus* C2A | 72% | 100% | 72% | 659.1 |
| *S. xylosus* DSM20266 | 72% | 100% | 72% | 931.6 |
| *S. equorum* LTH5015 | 74% | 98% | 72% | 635.8 |
| *S. epidermidis* E73 | 72% | 100% | 72% | 598.1 |
| *S. warneri* 197-70259724 | 73% | 99% | 73% | 1340.2 |
| *S. argenteus* DSM28299 | 92% | 100% | 92% | 425.1 |
| *S. schweitzeri* DSM28300 | 92% | 100% | 92% | 282.9 |
| *S. aureus* USA300 | 100% | 100% | 100% | 343.9 |

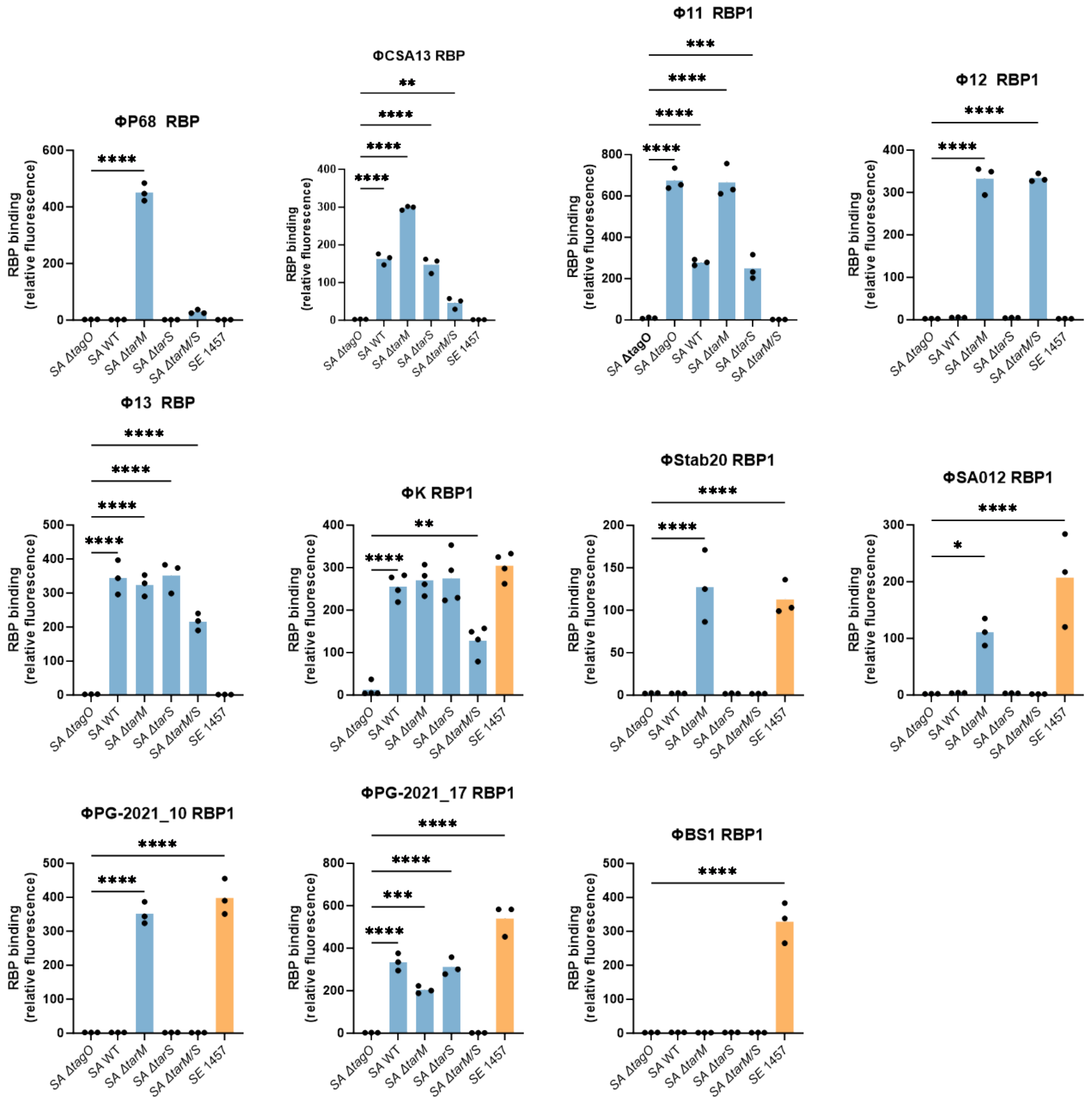


**Figure S1:** FACS-based assay data for RBPs. Median fluorescence of RBP treated bacteria. Significance calculated via ordinary one-way ANOVA and multiple comparisons were performed between the WTA-negative *S. aureus* Δ*tagO* strain and the differently glycosylated strains, *P < 0.05, **P < 0.01, ***P < 0.001, ****P < 0.0001.

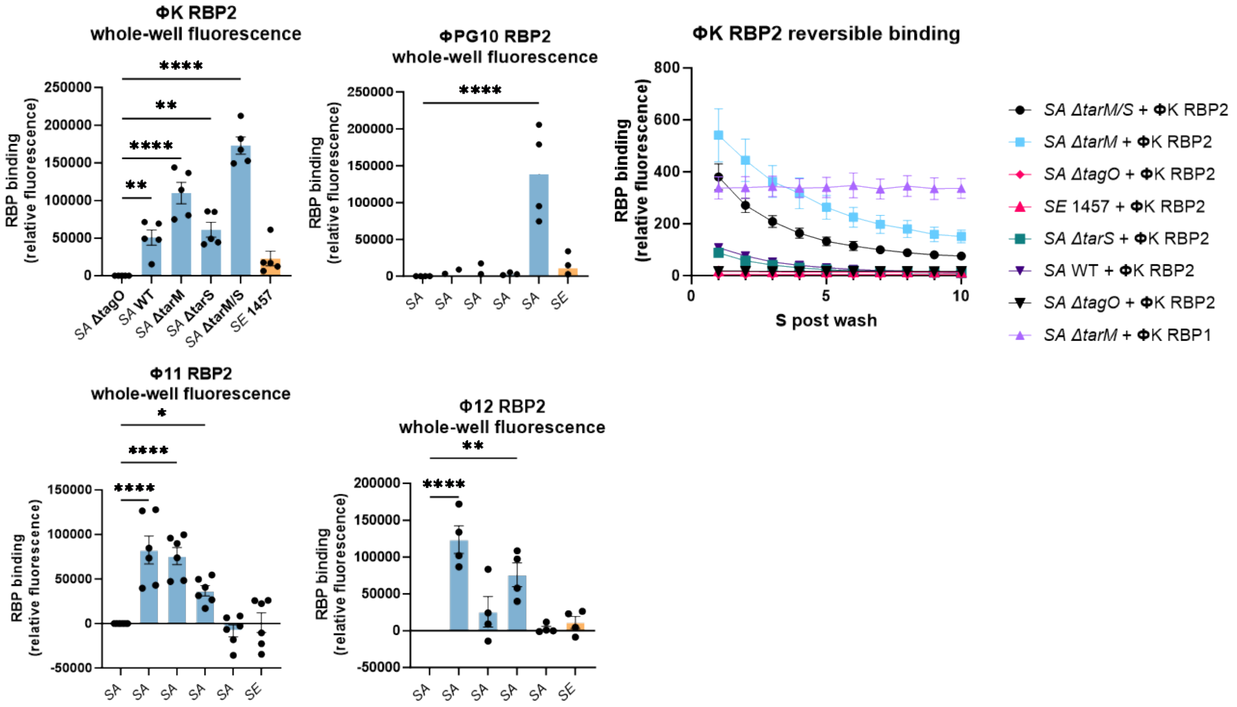


**Figure S2: a,** Fluorescence reader assay data of reversibly binding RBPs. Significance calculated via ordinary one-way ANOVA and multiple comparisons were performed between the WTA-negative *S. aureus* Δ*tagO* strain and the differently glycosylated strains, *P < 0.05, **P < 0.01, ***P < 0.001, ****P < 0.0001. **b,** Time-dependent dissociation of reversible ΦK RBP.

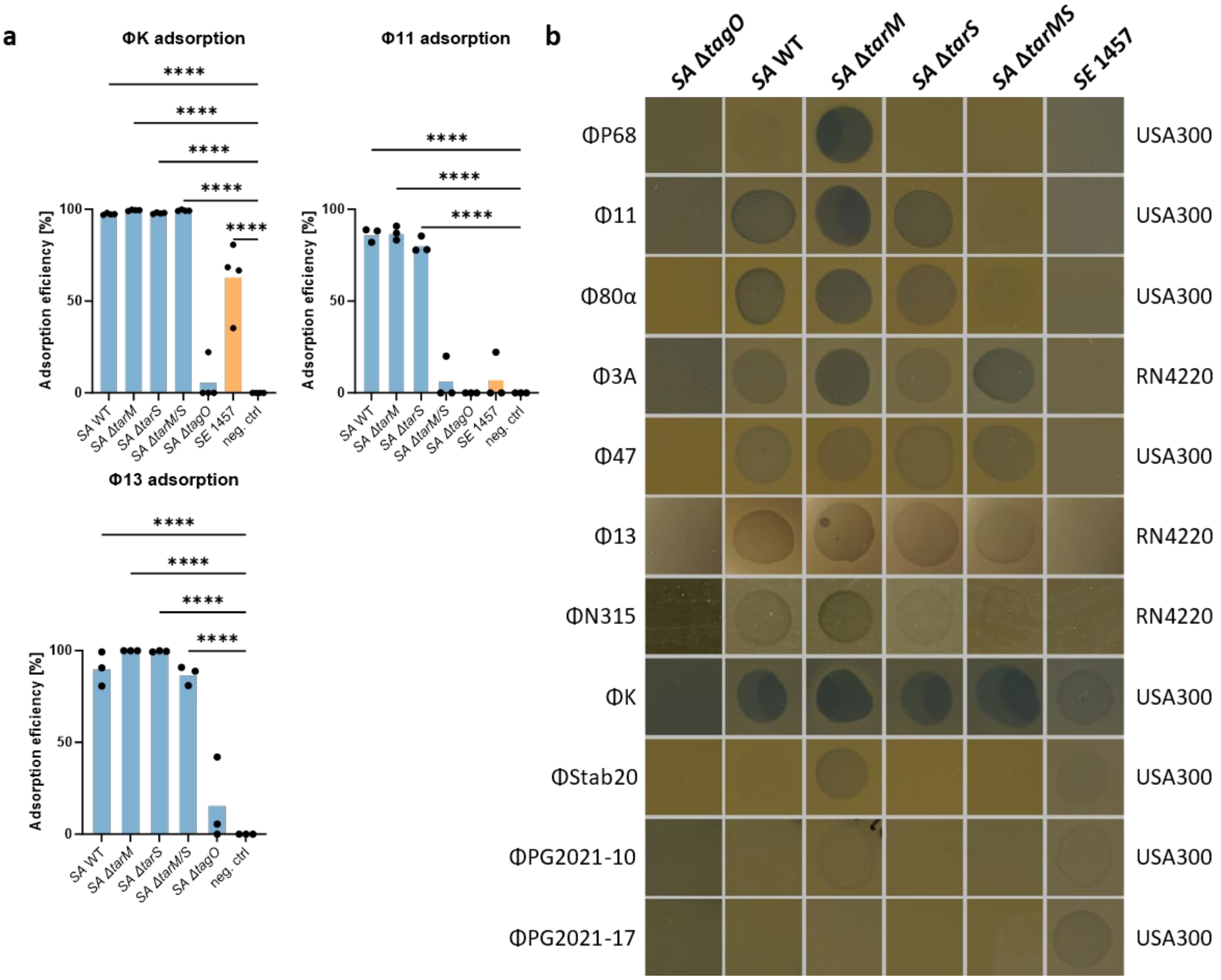


**Figure S3: a,** Adsorption assays of Φ11, Φ13, and ΦK. Significance calculated via ordinary one-way ANOVA and multiple comparisons were performed between the negative control and the different WTA strains, *P < 0.05, **P < 0.01, ***P < 0.001, ****P < 0.0001. **b,** Spot assays of various phages with appropriate phage dilutions.

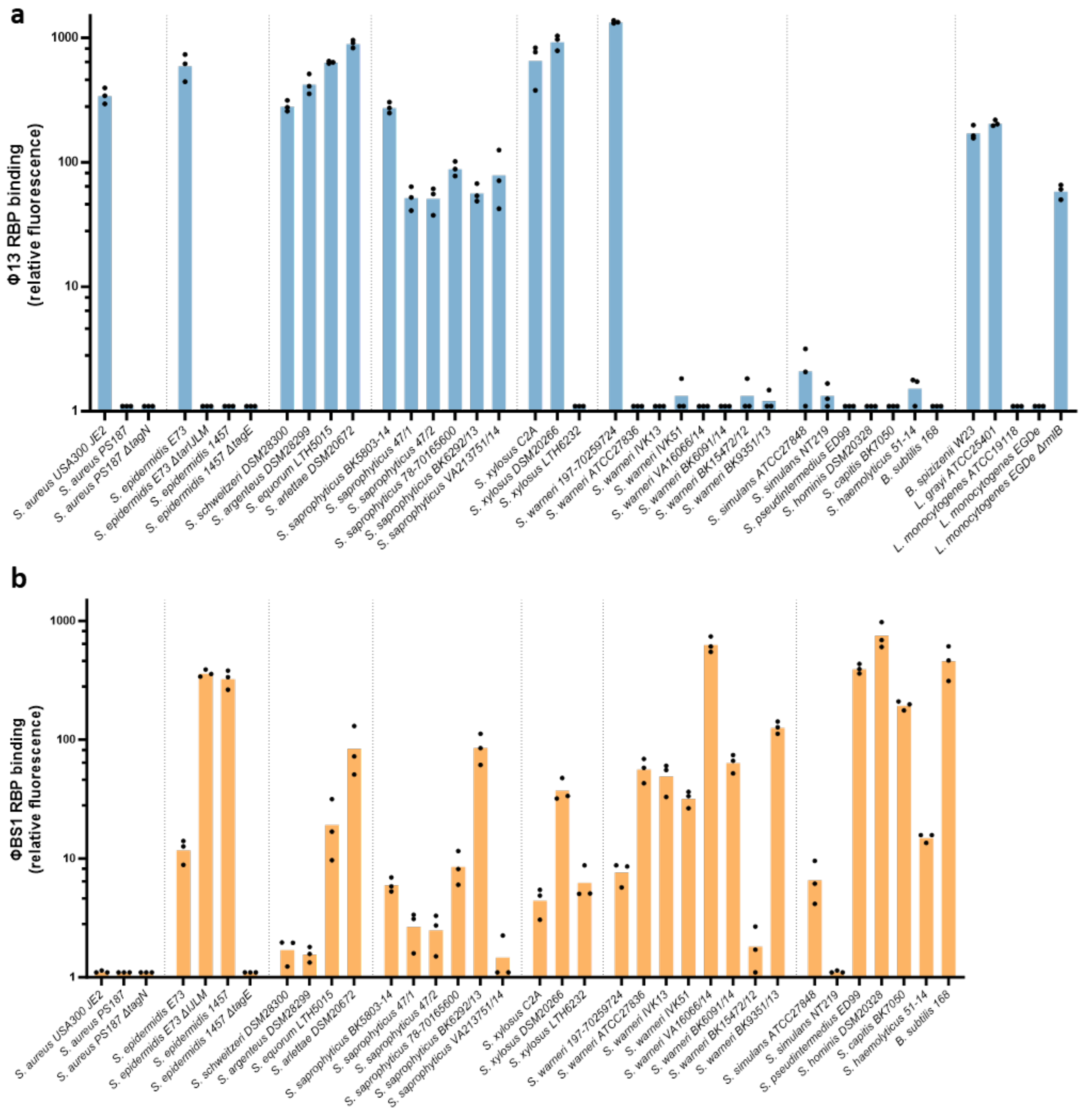


**Figure S4: a,** Φ13-RBP-binding signal of various bacteria. Φ13-RBP can only bind to RboP WTA. Genomic presence of the *tarIJL* cluster with similarity to USA300 *tarIJL* above 65% is indicated with +, absence is indicated with -. **b,** ΦBS1 RBP-binding signal of various bacteria. ΦBS1 can only bind to some GroP WTA glycosylation types. RboP can shield the GroP WTA and thus decrease binding of ΦBS1-RBP, as can be seen with *S. epidermidis* E73/*ΔtarIJLM*. Some bacteria showed strong signals with both, RboP (Φ13-RBP) and GroP (ΦBS1-RBP), such as *S. saprophyticus* BK6292/13. These strains could have both, RboP- and GroP WTA, that can be bound by different phages, potentially functioning as a hub for horizontal gene transfer between staphylococci with different WTA backbone structures.
